# Role of the prefrontal cortex in musical and verbal short-term memory: A functional near-infrared spectroscopy study

**DOI:** 10.1101/2023.10.19.563108

**Authors:** Jérémie Ginzburg, Anne Cheylus, Elise Collard, Laura Ferreri, Barbara Tillmann, Annie Moulin, Anne Caclin

**Author notes:** Corresponding author: Jérémie Ginzburg.

## Abstract

Auditory short-term memory (STM) is a key process in auditory cognition, with evidence for partly distinct networks subtending musical and verbal STM. The delayed matching-to-sample task (DMST) paradigm has been found suitable for comparing musical and verbal STM and for manipulating memory load. In this study, musical and verbal DMSTs were investigated with measures of activity in frontal areas with functional near-infrared spectroscopy (fNIRS): Experiment 1 compared musical and verbal DMSTs with a low-level perception task (that does not entail encoding, retention, or retrieval of information), to identify frontal regions involved in memory processes. Experiment 2 manipulated memory load for musical and verbal materials to uncover frontal brain regions showing parametric changes in activity with load and their potential differences between musical and verbal materials. A FIR model was used to deconvolute fNIRS signals across successive trials without making assumptions with respect to the shape of the hemodynamic response in a DMST. Results revealed the involvement of the dorso-lateral prefrontal cortex (dlPFC) and inferior frontal gyri (IFG), but not of the superior frontal gyri (SFG) in both experiments, in keeping with previously reported neuroimaging data (including fMRI). Experiment 2 demonstrated a parametric variation of activity with memory load in bilateral IFGs during the maintenance period, with opposite directions for musical and verbal materials. Activity in the IFGs increased with memory load for verbal sound sequences, in keeping with previous results with n-back tasks. The decreased activity with memory load observed with musical sequences is discussed in relation to previous research on auditory STM rehearsal strategies. This study highlights fNIRS as a promising tool for investigating musical and verbal STM not only for typical populations, but also for populations with developmental language disorders associated with functional alterations in auditory STM.

## Introduction

Auditory Short-Term Memory (STM) plays a crucial role in the processing of auditory information and enables understanding of a dynamic acoustic environment. STM is the active maintenance of information for a brief duration, typically lasting a few seconds (Baddeley & Hitch, 1974; Cowan, 2008). Auditory STM has been extensively studied for verbal material, but a growing body of research has also studied STM for musical material, revealing both shared and specific mechanisms depending on the type of sounds to maintain (Caclin & Tillmann, 2018). Domain-general mechanisms for processing time-based serial order in auditory sequences have been proposed (Gorin et al., 2016), while domain-specific systems have been suggested for encoding and maintenance of distinct materials (Schulze & Tillmann, 2013). Impairment of verbal STM has been shown to be a hallmark of language-related learning disorders, such as dyslexia (Majerus & Cowan, 2016; Roodenrys & Stokes, 2001; Ziegler et al., 2009, 2011) and Developmental Language Disorder (DLD, Archibald & Gathercole, 2006; Nickisch & Von Kries, 2009; Nithart et al., 2009). A few studies have also observed impairment for musical STM in dyslexic children (Couvignou et al., 2023; Forgeard et al., 2008; Ziegler et al., 2012). Moreover, recent studies have shown a sizeable comorbidity (∼30%) between dyslexia and congenital amusia (in both adults: Couvignou et al., 2019, and children: Couvignou & Kolinsky, 2021; Couvignou et al., 2023). Congenital amusia is a lifelong disorder characterized by a specific impairment in musical STM (for reviews, see Peretz, 2016; Tillmann et al., 2015, 2023). Overall, these findings suggest impairment of auditory STM in learning disorders, which could either be domain-general or domain-specific. Together with the various patterns of auditory STM impairments observed in neurological diseases (review in Caclin & Tillmann, 2018), they stress the importance of exploring auditory STM for both materials (music and speech).

### Fronto-temporal networks for musical and verbal STM

Neuroimaging studies of auditory STM have revealed the involvement of distributed brain networks including auditory areas in the superior temporal lobe, frontal areas, such as the dorso-lateral prefrontal cortex (dlPFC) and the inferior frontal gyrus (IFG), parietal areas including the supra-marginal gyrus, as well as other brain regions, notably the cerebellum, basal ganglia, and premotor areas (for a review, see Caclin & Tillmann, 2018). Only a few studies using functional magnetic resonance imaging (fMRI) investigated the brain networks involved in STM for both musical and verbal material. A first study reported overlapping brain networks in the superior temporal, inferior parietal, and frontal areas when participants rehearsed out loud novel melodies and nonsense sentences (Hickok et al., 2003). These findings were supported by a second study that examined sung syllables (Koelsch et al., 2009) in which the authors found similar networks activated during the silent rehearsal of verbal information (syllables) and tonal information (pitches), including mostly lateral frontal regions: left IFG and bilateral pre-motor areas, and a small cluster in the left planum temporale. A subsequent fMRI study with sung syllables compared musicians with nonmusicians and revealed that during encoding, parietal and temporal (auditory) cortices showed significant activation in both groups (Schulze et al., 2011). During rehearsal of verbal or tonal information, activation was observed in lateral prefrontal regions (including the dlPFC and IFG) and subcortical structures. While overlapping for both materials, activations were more extended in frontal areas for musical material in musicians. In a recent fMRI study, Albouy et al (2019) investigated STM for musical and verbal material using a delayed-matching-to-sample task (DMST) and compared activations with control perceptual tasks in amusic participants and matched control participants. During the maintenance of tonal and verbal information, activations emerged for control participants in left lateral frontal regions including the dlPFC and the IFG when comparing the memory task with the simple perception task. However, no activation was observed in temporal auditory regions, suggesting the specific involvement of lateral frontal regions for the maintenance of musical and verbal material, in line with previous findings (Hickok et al., 2003; Koelsch et al., 2009; Schulze et al., 2011). In contrast to controls, amusics’ brain activation patterns revealed several functional alterations during encoding and maintenance of musical information including in the right auditory cortex, the right dlPFC, and the right IFG. These results suggest that networks subtending musical and verbal STM are dissociable when studying a population with a specific deficit for the musical material. Overall, neuroimaging data support the view that temporal regions and lateral frontal regions are involved in the encoding of musical and verbal information, while lateral frontal regions are involved in the maintenance of these information, along with other parietal and subcortical areas in some studies. Moreover, these networks seem to be largely overlapping for both materials (Schulze & Koelsch, 2012) and evidence for specialized brain networks for musical and verbal STM arose in fMRI data only from the comparison of impaired (amusics) or expert (musicians) populations to controls.

### Memory load manipulation: n-back tasks

Investigating cognitive effort by manipulating memory load is valuable for understanding the underlying mechanisms of auditory STM. Most neuroimaging research on this topic has done so with the verbal identity n-back task that entails maintenance and manipulation of information, hence working memory (WM) (León-Domínguez et al., 2015). In this task, participants are presented with a series of stimuli (e.g., letters, digits) and are required to indicate when the current item matches the one presented n trials earlier. The task has the advantage of being applicable in the visual or auditory modality. In a meta-analysis of neuroimaging studies with verbal n-back tasks, Owen et al. (2005) found that, regardless of the modality, activity increases with memory load in frontal regions, including dlPFC, IFG, premotor cortex (PMC), and supplementary motor area (SMA). In a meta-analysis with n-back tasks, Sternberg tasks (Sternberg, 1966), and DMSTs, Rottschy et al. (2012) identified bilateral activation patterns where activity increased with memory load, encompassing frontal regions (dlPFC, IFG, PMC, SMA), the middle cingulate cortex and temporo-occipital areas, regardless of the task, modality, or material (words, pseudo-words, pictures…). Note that in this meta-analysis, all Sternberg tasks and DMSTs used visual material. Although the n-back task is frequently employed in neuroimaging studies to manipulate memory load, it is primarily designed to assess WM rather than STM. Indeed, the n-back task involves actively manipulating and updating information, which is specific of WM processes. Furthermore, neuroimaging studies have demonstrated that distinct lateral frontal regions are engaged in forward digit spans (STM) and backward digit spans (WM) (Owen, 2000; Tian et al., 2014). Therefore, the n-back task may not be the most suitable for specifically exploring the effects of memory load on STM per se, as it involves additional cognitive operations beyond simple storage, maintenance, and retrieval of information. Alternatively, recall tasks can be used to study STM and WM, but verbal serial recall tasks involve specific cognitive operation, as they depend on several phonological, sublexical, lexical and semantic factors (e.g. Allen & Hulme, 2006). Furthermore, serial recall tasks are not easy-to-use for studying musical material due to their reliance on production (Caclin & Tillmann, 2018; but see Gorin et al., 2018; Williamson et al., 2010 for mixed recall/recognition task adaptations to music). Therefore, DMSTs seem to be specifically suited to investigate memory load in auditory STM. However, they have only rarely been employed in neuroimaging studies manipulating memory load in auditory STM.

### Memory load manipulation: DMST

Auditory DMST consists in making a same/different judgement between two sequences separated by a silent delay (Albouy et al., 2019; Caclin & Tillmann, 2018; Ginzburg et al., 2022; Gosselin et al., 2009; Talamini et al., 2021; Tillmann et al., 2009). This paradigm lends itself to memory load manipulation for musical and verbal material, to study both STM and WM (with forward and backward instructions respectively), and has been used with three types of auditory material in behavioral experiments (words, pitch, and timbre sequences: Schulze et al., 2012; Schulze & Tillmann, 2013). Research using electroencephalography (EEG) with auditory DMST has consistently demonstrated that a sustained anterior negativity (SAN) in the electrophysiological signal serves as a reliable index of memory load. As memory load increases, the amplitude of SAN at fronto-central electrodes increases for both pitch and timbre materials (Alunni-Menichini et al., 2014; Guimond et al., 2011; Lefebvre et al., 2013; Lefebvre & Jolicœur, 2016; Nolden et al., 2013). However, due to the limited spatial resolution of EEG, the precise engagement of frontal regions could not be fully explored. Only one neuroimaging study, using magnetoencephalography (MEG), has thus far investigated the specific brain regions showing parametric activation with memory load increase during DMST with musical material (Grimault et al., 2014). By examining MEG data at the end of the silent retention delay, significant correlations between the memory load and the amplitude of the evoked response were reported for bilateral IFGs, bilateral dlPFCs, bilateral auditory cortices, and the right parahippocampal gyrus. These results indicate that lateral frontal regions (IFG and dlPFC), previously identified to be associated with musical and verbal STM (Albouy et al., 2019; Koelsch et al., 2009; Schulze et al., 2011), show a parametric increase of activity with musical memory load, in keeping with neuroimaging studies using verbal n-back tasks (Owen et al., 2005; Rottschy et al., 2012).

### fNIRS studies with memory load manipulation in n-back tasks

In recent years, there has been a growing interest for using functional Near-Infrared Spectroscopy (fNIRS) to study brain functioning. fNIRS is a neuroimaging technique that utilizes light sources and detectors at near-infrared wavelengths to measure changes in cerebral metabolism, which serve as an indirect measure of neuronal activity. When neuronal activity increases in a particular cortical area, a rise in oxygenated hemoglobin (HbO) and a decrease in deoxygenated hemoglobin (HbR) occur concurrently, which can both be recorded with fNIRS. This hemodynamic response peaks around 5 seconds after stimulus onset (Fantini et al., 2018; Huppert et al., 2006; Scholkmann et al., 2014). The temporal dynamics of the fNIRS signal is thus similar to the blood-oxygen level-dependent (BOLD) signal in fMRI. Although having a physiological delay typical of hemodynamic signals and a limited spatial resolution, including the difficulty to record deep brain areas (such as the auditory cortex), fNIRS offers several advantages over other imaging techniques, such as fMRI (Pinti et al., 2020). It provides a quiet and less restrictive environment as it is non-invasive and portable, making it more comfortable and suitable for capturing real-world behavior and associated neural responses in children and clinical populations who may find the fMRI environment distressing (Ferreri et al., 2014). Additionally, fNIRS has a higher tolerance to movement compared to fMRI and electroencephalography (Aslin & Mehler, 2005). Furthermore, whereas the loud sounds of fMRI can present challenges for studying the auditory modality (and in particular for individuals with language processing difficulties as it requires them to listen in a noisy environment), fNIRS is fully silent (Butler et al., 2020; Hancock et al., 2023).

A number of fNIRS studies have employed verbal n-back task to investigate memory load. In n-back tasks with auditorily or visually presented consonants, activity in bilateral lateral prefrontal cortices (LPFCs, including dlPFC and IFG) increased with increasing memory load (Rovetti et al., 2021), while bilateral medial prefrontal cortices (MPFCs) do not seem to respond to memory load manipulation. These results are in accordance with previous neuroimaging studies showing the involvement of LPFC structures in WM, with activity increasing parametrically with load (Owen et al., 2005; Rottschy et al., 2012). Using major, minor, and dissonant chords as stimuli in a n-back task, Tseng et al. (2018) showed that activity increased parametrically with load in bilateral orbital PFC and IFG. However, the use of complex chords (more than three tones simultaneously) and major/minor/dissonant chords possibly entailed that participants categorized these stimuli based on the abstract representation of consonance/dissonance, rather than pure pitch information. Recording the same brain regions as in Rovetti et al (2021), hearing-aid users and normal-hearing participants displayed an increase of activity with load, leveling at 2-back difficulty in ventral and medial PFCs (Rovetti et al., 2019). This parametric activation was observed for verbal memory load in the auditory and visual modality. To the best of our knowledge, only one study used fNIRS to investigate auditory WM load in children with developmental language disorder (DLD) and typically developing children (TD) using an auditory n-back task with consonants (Hancock et al., 2023). There was an increase of activity with increasing memory load in the left dlPFC (the right dlPFC was not recorded) and a decrease of activity with increasing memory load in bilateral inferior parietal lobules (IPLs) in the TD group, but not in the DLD group. These results suggest a relationship between DLD and difficulties in engaging neural activity for different auditory WM load in dlPFC and parietal regions. In summary, similarly to fMRI investigations, fNIRS studies using auditory n-back tasks have consistently shown the involvement of lateral frontal regions (IFG and dlPFC) in STM, with activity increasing parametrically with load. In contrast, medial frontal areas appear to have a limited involvement in STM.

### Studying auditory STM with Delayed-Matching-to-Sample Tasks in fNIRS

One methodological study has shown that DMST is a suitable paradigm to investigate auditory STM using fNIRS, using only verbal material and a single sequence length (Yamazaki et al., 2020). After listening to (or watching at) a 9-syllable sequence, participants had to maintain the information during a 9-seconds retention delay and compare it to a second 9-syllable sequence that could be either identical or different by one syllable. Within a large array of recording channels over the left frontal and temporal areas, significant activation during the encoding and maintenance phase was observed in the auditory modality in the left IFG and dlPFC respectively, along with other premotor and temporal areas.

The primary objective of the current study was to investigate auditory STM, and more specifically the effect of memory load, for musical and verbal material using a DMST paradigm. Two experiments were conducted to achieve this goal. For both experiments, our focus was on the frontal brain regions consistently reported to be involved in both musical and verbal STM (i.e., IFG and dlPFC). Auditory cortices were not targeted with fNIRS due to their depth in location. Additionally, we recorded medial frontal regions (SFG) as control, since these areas are not typically activated with the fronto-temporal network involved in auditory STM. Experiment 1 adapted the experimental design of Albouy et al. (2019) comparing musical and verbal DMST to a low-level perception task of equivalent duration. Our aim was to replicate findings obtained with fMRI, namely that the lateral prefrontal regions exhibited stronger activation during the memory task compared to the low-level perception task. In Experiment 2, memory load for musical and verbal material was manipulated by varying sequence length. Overall, the research questions were as follow: (1) Is fNIRS suited to explore frontal activations during auditory DMST? (2) What are the frontal brain regions where activity varies parametrically with memory load? (3) Are these regions similar for musical and verbal material?

## Experiment 1

### Methods

#### Participants

Nineteen healthy adults were recruited for Experiment 1. Data from three participants were excluded because of technical problems in the fNIRS signal acquisition. This led to a final sample of sixteen right-handed participants (mean age = 39.2 years, sd = 15.2 years, min = 21 years, max = 62 years, 12 females, mean education level = 14.95 years). They all gave written informed consent to participate in the experiment. Prior to the main experiment, all participants were tested with pure tone audiometry (separately for the two ears, 250 Hz, 500 Hz, 1000 Hz, 2000 Hz, 4000 Hz, 6000 Hz, 8000 Hz), the Montreal Battery of Evaluation of Amusia (MBEA, Peretz et al., 2003) and a Pitch Discrimination Threshold (PDT) test (Tillmann et al., 2009). All participants presented a normal audiometry (hearing threshold lower than 30 dB at any frequency in both ears). No participant presented any pitch perception or pitch memory impairment (MBEA > 25 (maximum score = 30) and PDT < 1 semitone) and they had no or little musical education (mean musical education = 0.1 years sd = 0.5 years). No participant presented any neurological or psychiatric history and none reported any past diagnosis of neurodevelopmental disorder. All study procedures were approved by a national ethics committee (CPP Ile de France VI, ID RDC 2018-A02670-55) and participants received a compensation for their participation. During the first session (∼1h30), participants performed the audiometry, MBEA, and PDT. During the second session (∼1h15) on a different day, participants underwent the fNIRS testing (see below).

### Stimuli construction and task design

#### Musical and verbal stimuli

Musical and verbal stimuli were the same as in Ginzburg et al. (2022). For musical tasks, six musical tones (created with the software Cubase 5.1 (Steinberg) and a Halion Sampler (Steinberg) using an acoustic piano timbre) belonging to the C major scale were used (C2, E2, G2, B2, D3, F3) with frequencies ranging from 131 to 349 Hz (thus encompassing the fundamental frequency range of the vowel recordings: 202-212 Hz). For verbal tasks, the items were Consonant-Vowel syllables that were selected to show the greatest perceptual distance with each other. Six consonants and six vowels were selected: /f/ /t/ /z/ /g/ /m/ /l/ and /i/ /e/ /a/ /y/ /ø/ /u/, thus resulting into 36 syllables that were recorded by a professional mezzo-soprano singer (for details about syllables construction, see Supplementary Figure S1 in Ginzburg et al., 2022). Six syllables were selected: /fi/ /gu/ /ly/ /mø/ /te/ /za/.

Trial examples can be found at https://github.com/jeremieginzburg/supp_mat_STM_adults_fNIRS.

#### Perception and memory tasks

For each trial of perception and memory tasks, participants were asked to listen to two 5-item auditory sequences (S1 and S2; verbal or musical) separated by a silent retention delay of 6000 ms. Each item lasted 500 ms, the silent inter-stimulus interval (ISI) between two items lasted 100 ms. Overall, there was a 600 ms stimulus onset asynchrony (SOA), leading to a duration of 11800 ms for S1-delay-S2. Participants were given 3000 ms to provide a same/different response after the end of S2. The next trial started after a 5000-to 9000-ms randomly-jittered silent interval after the response window. Presentation® software (Version 18.0, Neurobehavioral Systems, Inc., Berkeley, CA, www.neurobs.com) was used to present stimuli and record responses. Additionally, sixteen 22800-to 26800-ms randomly-jittered silent trials were generated to intersperse within each testing block (see *Procedure* below).

For the memory task, S1 and S2 sequences could be either identical or different. All items were different within a given sequence, and all musical sequences included at least one ascending interval and one descending interval. When S2 was different, a new item could appear equiprobably at the 2^nd^, 3^rd^, 4^th^ position (positions 1 and 5 were not used for changes to minimize primacy and recency effects). For each material, six S1 sequences were used to create “same” trials, with S2 identical to S1. Six other S1 sequences were used to create “different” trials, with S2 sequences differing from S1 by one item. For the musical material, the new item in S2 always changed the contour of the sequence (the contour is the up-and-down scheme of a melody). So, if S1 had a down-up-**down**-up contour (e.g. E2-C2-**B2-<u>G2</u>**-F3), S2 could have a down-up-**up**-up contour (e.g. E2-C2-**B2-<u>D3</u>**-F3).

For the perception task, new sequences were created and S1 and S2 were always different. In the S2 sequence, the last two items could either be identical or different. Except when S2 contained two identical last items, all items were different within a given sequence, and for the musical material all sequences included at least one ascending interval and one descending interval. For the musical material, when the last two items were different, these two items could not differ by more than 3 tones. A total of twelve S1-S2 sequences were created for each material, half of them with S2 sequences having two identical last items and half of them with S2 sequences having two different last items. For this perception task, the participants were asked to ignore S1 and to answer whether the two last items of S2 were the same or different.

#### fNIRS montage and data acquisition

The absorption of near-infrared light was measured at 760– and 850-nm wavelengths at a sampling frequency of 7.81 Hz using a continuous-wave NIRScout device (NIRx Medical Technologies, LLC). The data were collected using the NIRStar 15.3 acquisition software. Eight light sources and twelve light detectors were attached to a cap with a 10-20-system marking for probe placement. Additionally, eight 8-mm short-distance channels (one for each source) recorded systemic signal.

The montage was created using fOLD (fNIRS Optodes’ Location Decider, Zimeo Morais et al., 2018), which allows placement of optodes in the international 10-20 system to maximize coverage of chosen anatomical regions as defined by one of five segmentation atlases. For the segmentation atlas, we chose the AAL2 (Automated Anatomical Labeling, Rolls et al., 2015) to generate a montage (Figure 1) covering the inferior frontal gyri (IFG) and dorsolateral prefrontal cortex (dlPFC) as they were shown with fMRI to be involved in the encoding and retention phase in memory recognition tasks (Albouy et al., 2019). As a control, our montage additionally covered medially the superior frontal gyri (SFG), as they have not been shown to be involved in memory recognition tasks.

**Figure 1:**
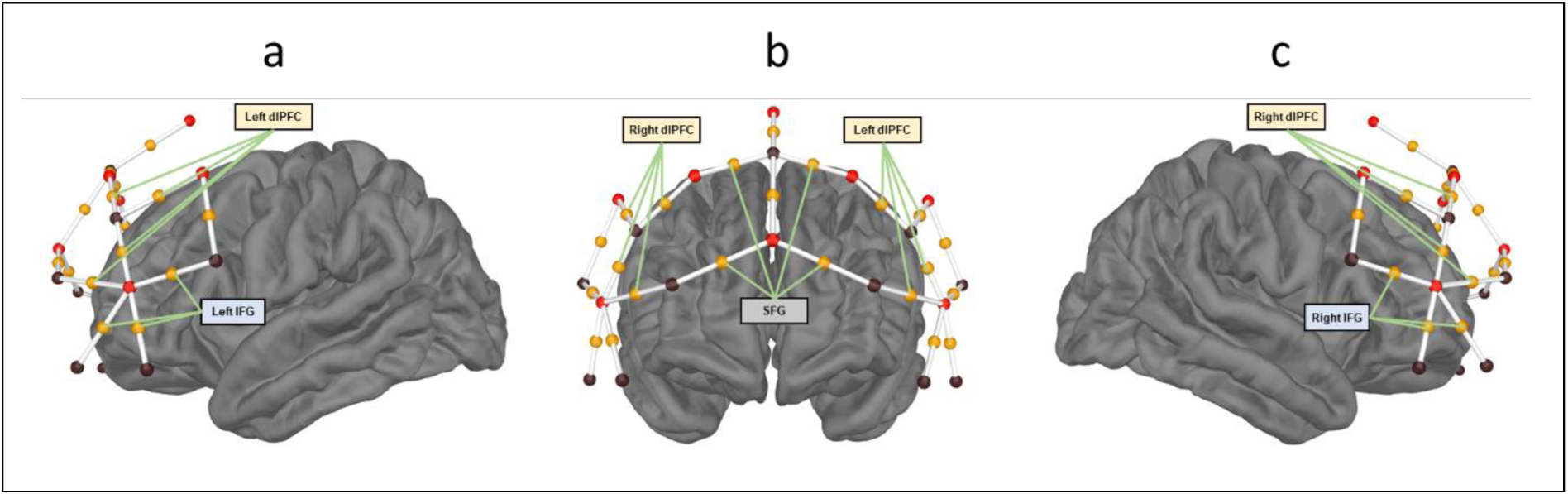
Location of sources (black spheres) and detectors (red spheres) and their corresponding midpoint channels (white lines with orange spheres) for (a) left; (b) front; and (c) right views of the brain. Green lines illustrate the ROIs used for analysis: 3 channels for each IFG (lateral views), 4 channels for each dlPFC (front and lateral views), 4 channels covering bilateral SFGs (front view).

The final montage included 22 measurement channels with distances ranging from 2.9 to 4.1 centimeters (the montage is represented on Figure 1 and see Supplementary Table 1 for maximum recording specificity of each channel). Data from two channels that had the highest specificity for the precentral gyrus were not analyzed as they were not in the scope of the current study. We discarded two midline channels because they presented noisy signal. Overall, we retained 18 channels for analysis. Among those channels and as calculated by the fOLD software, in each hemisphere, three of them had the highest level of recording specificity for IFG and four of them for dlPFC. Four channels had the highest specificity for bilateral SFGs.

#### Procedure

The experimenter measured the participant’s head circumference to determine cap size. Cap alignment was verified and adjusted if needed so that the probe at Cz was located halfway in the nasion-to-inion and the tragus-to-tragus measurements. The participant was then led into a dimly lit, sound-attenuated booth where, if needed, the participant’s hair was moved around the optode locations using a thin wooden stick to provide clear access to the scalp. Optodes were then placed according to the pre-established montage. fNIRS signal was calibrated, checked for quality, and optode placement on the scalp was readjusted until a satisfying signal quality was obtained before proceeding. Right before testing, the experimenter gave the task instructions to the participant.

Presentation of stimuli and recording of behavioral responses was controlled by Presentation^©^ software (Version 18.0, Neurobehavioral Systems, Inc., Berkeley, CA, www.neurobs.com). Triggers were sent from Presentation^©^ to the fNIRS acquisition system using a parallel port. Participants sat approximately 50 cm from the computer screen, auditory stimuli were played through Sennheiser HD-250Pro headphones at a comfortable intensity (80 dB SPL A-weighted measured with a Larson-Davis System 824 with an AEC101 IEC 318 Artificial Ear Coupler). Headphones were positioned behind the optodes (primarily located on frontal area) and were sufficiently flexible to avoid any interference with the optical fibers. Participants gave their response with a computer mouse with their right-hand (left-click for ‘same’ response and right-click for ‘different’ response).

All participants underwent four blocks of sixteen trials, one block per task and per material: musical perception, musical memory, verbal perception, verbal memory. Among the sixteen trials, twelve of them were stimulation trials as described above. Half of the trials were “same” trials and half of the trials were “different” trials. Stimulation trials were pseudo-randomly presented with the constraint that no more than three consecutive ‘same’ (or ‘different’) trials could be presented. Four “silent” trials were played on the 1^st^, 6^th^, 11^th^, and 16^th^ trial position of a block in order to allow for the hemodynamic signal to return to baseline (Balters et al., 2021) and to serve as an implicit baseline in the Finite Impulse Response (FIR, see below) analysis (Cairo et al., 2004; Kharitonova et al., 2015). Each of the four blocks lasted around 6 minutes and was preceded by a four-trial training block with feedback using the same task and material. For the test blocks, no feedback was given. The order of test blocks was counterbalanced across participants using a Latin square balancing for first-order carryover effects using the *crossdes* R package (Sailer, 2022).

#### Behavioral data analysis

Measures of d-prime (d’) and criterion (c) were obtained according to Signal Detection Theory (SDT) for each task, material, and participant (Macmillan & Creelman, 2005). Hit corresponded to a correct answer for a different trial. False alarm corresponded to an incorrect answer for a same trial. *d’*, or sensitivity, was calculated using the *psycho* package (Makowski, 2018) as the z-score of False Alarms subtracted from the z-score of Hits. The criterion was calculated as the mean z-score of Hits and False-alarm rates multiplied by minus one and reflect an observer’s bias to say yes (in our case “different”) or no (“same”), an unbiased observer having a value around 0. A liberal bias (tendency to say “different”) results in a negative c, a conservative one results in positive c. Correction of extreme values was made following Hautus (1995) who recommends the use of a log-linear rule that consists of increasing each cell frequency of the contingency table by 0.5, irrespective of the content of each cell. Furthermore, we analyzed the response times (RT) of participants after the end of S2 for correct trials only, averaged separately for each task, material, and each type of trial (same/different). Note that participants had 3 seconds to answer, otherwise the response was counted as irrelevant and thus not considered in the behavioral analysis. Those missed trials represented 0.01 % of all trials.

Analyses were conducted using Bayesian Statistics that allow the direct comparison of the predictions of several hypotheses (including the null model) and to estimate a degree of logical support or belief regarding effects of interest and their interactions (Wagenmakers et al., 2018). We report Bayes Factor (BF_10_) as a relative measure of evidence of an effect compared to the null model. Traditionally, a BF_10_ between 1 and 3 is considered as weak evidence for the tested model, between 3 and 10 as positive evidence, between 10 and 100 as strong evidence and higher than 100 as decisive evidence. Similarly, to interpret the strength of evidence in favor of the null model, a BF_10_ between 0.33 and 1 is considered as weak evidence, a BF between 0.01 and 0.33 as positive evidence, a BF between 0.001 and 0.01 as strong evidence and a BF lower than 0.001 as decisive evidence (Lee & Wagenmakers, 2014). For clarity purposes, we report information about the best model only.

Using the R *BayesFactor* package (Morey & Rouder, 2022), d’ and c were submitted to a Bayesian repeated-measure ANOVA including task (two levels: perception and memory), material (two levels: musical and verbal), and their interaction as fixed factors. Overall, four models were tested (task, material, task + material, task + material + task:material) and compared to the null model. As recommended by Van Den Bergh et al. (2022), participants were added to all models as random factors using the *lmBF* function of the *BayesFactor* package. Paired Bayesian t-tests were performed as post-hoc tests if the best model included the interaction. Correct RTs were submitted to the same Bayesian ANOVA, with the addition of the type of trial (same/different) factor.

Additionally, one-sample Bayesian t-tests against 0 were performed on the criterion for each task and material.

Furthermore, we report the results of the analysis of effects using the *bayesfactor_inclusion* function from the R from the *bayestestR* package (Makowski et al., 2019) that compares between models that do or do not incorporate a specific effect, such as a factor or an interaction. The resulting measure, BF_inclusion_, serves as a relative indicator of the evidence favoring the inclusion of a factor.

#### fNIRS data pre-processing and signal deconvolution

fNIRS data were pre-processed using the NIRS Brain-AnalyzIR toolbox (Santosa et al., 2018) and custom-written scripts. Raw intensity signals were first converted to changes in optical density. To correct for motion artifacts from excessive head movements, we applied Temporal Derivative Distribution Repair (TDDR) (Fishburn et al., 2019), a robust regression approach to remove large fluctuations in the optical density signal (motion artifacts), while keeping smaller fluctuations (hemodynamic activity). Corrected optical density were then band-pass filtered between 0.01 and 0.2 Hz to remove cardiac (∼ 1.2 Hz) and respiratory activity (∼ 0.25 Hz). Finally, corrected and filtered optical densities were transformed into (de)oxygenated hemoglobin concentrations using the modified Beer-Lambert Law.

Data were then processed with a General Linear Model (GLM) using a Finite Impulse Response (FIR) model to deconvolute the signals from successive trials for each channel. This method does not make assumptions regarding the shape of the hemodynamic response and allows for an unconstrained estimation of the full hemodynamic response during stimulation and maintenance. To do so, 36 one-second boxcar regressors were fitted around S1 onsets to encompass the total duration of stimulation (–5 seconds to 30 seconds around S1 onset) for each task and material. A boxcar regressor per block (encompassing the entire twelve stimulation trials and four silence trials) was added to account for possible HbO/HbR signal changes across blocks of recordings. Finally, data from all short-channels (eight HbO and eight HbR measures) were orthogonalized and added as regressors of no-interest in the GLM in order to further clean the signal from systemic components (Luke et al., 2021). Overall, for each chromophore (HbO/HbR), each recording channel of each participant was regressed using an Ordinary Least Squares (OLS) GLM with a design matrix including block regressors, 36 1-second boxcar regressors for each task and material (thus 144 regressors for the FIR models), and all short-channel signal (16 regressors). Note that no baseline correction is needed in the FIR GLM approach as silent trials, for which no regressors are fitted in the GLM, act as implicit baseline (Cairo et al., 2004; Kharitonova et al., 2015).

#### fNIRS data analyses

Deconvoluted data were then analyzed using a Bayesian ANOVA on beta coefficients for each 1-s time window of the FIR models and each ROI (see below). The use of Bayes factors to analyze time-course data shows great promise with robustness to type I errors without the need for corrections (see Teichmann et al., 2021 for an empirical comparison between cluster-based corrected time-course data and the Bayes Factor approach). Moreover, as compared to the traditional frequentist approach to time-course data that usually only allows for the comparison of two conditions when applying corrections for multiple testing, Bayesian statistics allow testing for multiple factors and their interaction at each time sample.

Five ROIs (see Figure 1 and Supplemental Table 1) were created with the channels showing the largest specificity for the left or right IFG (3 channels in each hemisphere), the left or right dlPFC (4 channels in each hemisphere), the bilateral SFGs (4 channels). For each participant, time point, task, and material, betas were averaged across channels making up each ROI. These ROI-averaged betas were then tested across participants, for each time point from –5 to 18 seconds around S1 onset (i.e., until 6 seconds after the end of S2, as the deconvoluted signal would then include the motor response which we did not intend to analyze) for each ROI with a repeated measure Bayesian ANOVA, as for behavioral data, including task (two levels: perception and memory), material (two levels: musical and verbal), and their interaction as fixed factors. Participants and their interaction with the task and material factors (random slopes) were added to all models as random factors. We report here the best model (as compared to the null model, BF_10_ > 1) for each time sample. Paired Bayesian t-tests were performed as post-hoc tests if the best model included the interaction.

We report in the main text only HbO results, as HbO tends to show higher amplitude changes and a higher SNR (Pinti et al., 2020) than HbR. However, HbR results are available in supplementary materials and commented in the discussion.

## Results

### Behavioral results

For d’ (Figure 2b), the best model explaining the data included both fixed factors (task and material) and their interaction (BF_10_ = 3.5e+5). The analysis of effects across matched models revealed decisive evidence for the task (BF_inclusion_ = 9.76e+03) and the material (BF_inclusion_ = 278.57) effects and weak evidence for the task:material interaction (BF_inclusion_ = 1.06). d’ was higher for the verbal material as compared to the musical material and higher for the perception task as compared to the memory task (Figure 2b). Post-hoc Bayesian t-tests revealed strong evidence for the task effect in the musical material (BF_10_ = 39.46) and positive evidence for the task effect in the verbal material (BF_10_ = 8.33).

**Figure 2:**
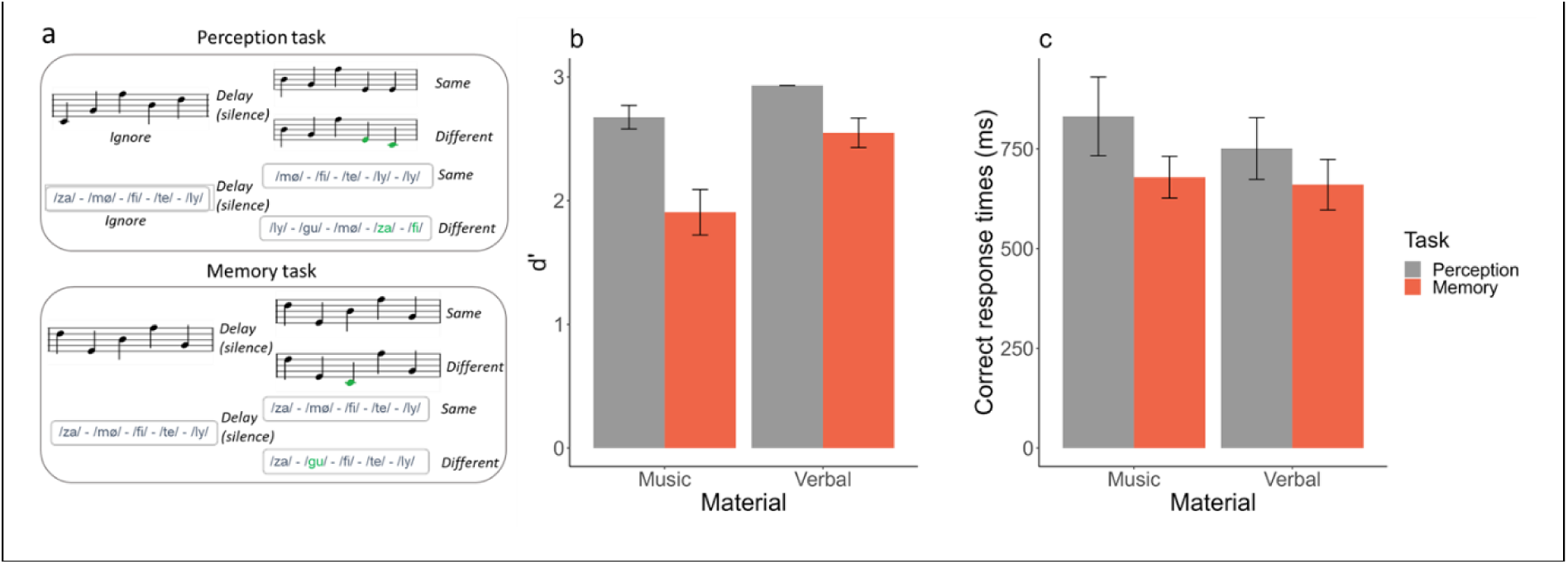
Tasks and average performance as a function of task (perception in grey /memory in orange) and material (musical/verbal). (a) Examples of trials for the perception and memory tasks for musical and verbal materials. (b) Mean and standard error of participant’s sensitivity (d’). (c) Mean and standard error of averaged response times for correct trials (time in millisecond that participants spent after the end of S2 before giving a “same” or “different” answer).

For correct RTs (Figure 2c), the best model was the one including only the task effect, but with only weak evidence relative to the null model (BF_10_ = 2.8). RTs were slightly longer in the Perception task compared to the Memory task.

For c, results are reported in supplementary Figure S1.

#### fNIRS results

HbO deconvoluted signals within the targeted ROIs are represented with corresponding statistics in Figure 3 (topographic representations are available in supplementary Figure S2 and HbR results in supplementary Figure S3).

**Figure 3:**
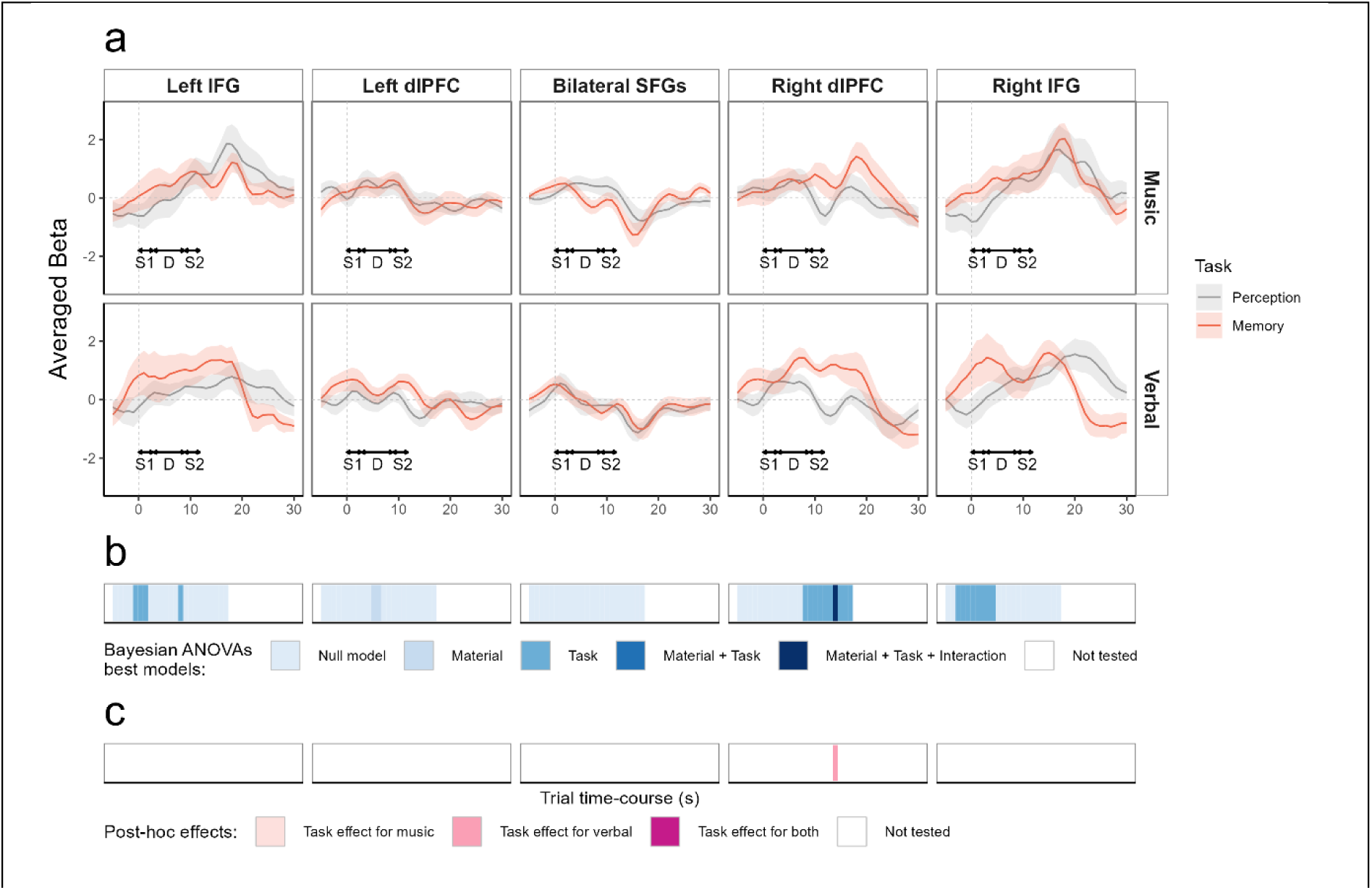
(a) Average beta (plain line) and standard error (shaded area) from the FIR deconvolution performed on HbO data for all participants (n=16), across five ROIs (left and right IFG, left and right dlPFC, bilateral SFGs), in a time window ranging from –5 to 30 seconds around S1 onset (grey dotted vertical line), for the perception task (grey) and the memory task (red). Top panel: musical material; bottom panel: verbal material. For clarity purposes, S1, D: silent retention delay, and S2 durations are indicated with double-headed arrows. (b) Time-course representation of Bayesian analysis, each blue shade represents the best model as compared to the null model (BF_10_ > 1). No statistics were performed after 18 seconds (end of S2 + peak of the hemodynamic response, ∼ 6s), because the deconvoluted signal would then include the motor response which we did not intend to analyze. (c) time-course representation of Bayesian post-hoc analysis performed only if the best model included the interaction. Purple shades indicate if there was evidence in favor of the task effect (BF_10_ > 1) for the musical material, the verbal material, or both.

For the left IFG, we found weak evidence from –1 to 1 seconds around S1 onset for the model including the task effect (1.47 < BF_10_ < 1.84) and weak evidence at 8 seconds for the model with the task effect (BF_10_ = 1.03), with in both cases higher betas for the memory task than for the perception task.

For the right IFG, we found weak to positive evidence from –3 to 4 seconds around S1 onset for the model including the task effect (1.11 < BF_10_ < 7) with higher betas for the memory task than for the perception task.

For the left dlPFC, we only found weak evidence at 5 and 6 seconds for the model including the material effect (1 < BF_10_ < 1.08), with higher betas for the musical material than for the verbal material.

For the right dlPFC, we found weak to decisive evidence for the model including the task effect from 8 to 17 seconds around S1 onset (1.07 < BF_10_ < 2.9e+03) with higher betas for the memory task than the perception task. At the 14 second post-S1 time sample, we found strong evidence for the model including all effects and their interaction (BF_10_ = 56.5). Post-hoc tests at this time sample revealed strong evidence for the task effect only for the verbal material (BF_10_ = 27) with higher betas for the memory task than the perception task.

For the SFG, no model was better than the null model to explain the data (all BF_10_ < 0.76).

## Discussion

Experiment 1 showed better behavioral performance and faster response times for the verbal material, as compared to the musical material for both perception and memory tasks. These results are in accordance with a previous study testing children with the same stimuli and the same memory paradigm that reported higher performance for the verbal material than for the musical material (Ginzburg et al., 2022). Other studies using different verbal stimuli, but the same memory task, reported a material effect in the opposite direction as observed here (verbal performance < musical performance, Albouy et al., 2019; Tillmann et al., 2009). This is due to the use of other verbal stimuli than the current study: mono-syllabic words, resynthesized to obtain a constant fundamental frequency and differing only by consonants in order to exploit the phonological similarity effect (Baddeley, 1966). In contrast, our verbal material aimed for phonologically distinct items as they differ in vowel as well (hence, easier perception and more contrasting material to be memorized). As expected and as in Albouy et al (2019), we also observed higher performance in the perception task compared to the memory task for both materials. Analysis of criterion revealed a weak tendency for a conservative bias for the musical material (more errors in ‘different’ trials) and a more liberal bias for the verbal material (more errors in ‘same’ trials). Based on these results, we increased the sequence length of the verbal material as compared to the musical material for Experiment 2, in order to equalize performance between materials.

HbO deconvoluted fNIRS signals over frontal areas showed a clear pattern of activation in lateral frontal channels over the time-course of the trial for both materials and both tasks (see supplementary Figure S2), with a higher increase in activity for the memory tasks as compared to the perception tasks. Conversely, a deactivation takes place in channels placed above medial frontal areas but does not seem to be different for the memory and the perception tasks. Analyses of activation in frontal ROIs showed evidence for higher activation in the memory task as compared to the perception task in the right dlPFC for both materials between 8 and 17 seconds after S1 onset (BF_10_ > 1) and in particular 10 to 13 seconds after S1 onset (all BF_10_ > 107). Considering that the hemodynamic response peaks ∼5-6 seconds after stimulus onset (Pinti et al., 2020), signals for these time samples should mainly correspond to the cortical activity during the silent retention delay (2.9 – 8.9 s). In bilateral IFGs, we also observed higher activation in the memory task compared to the perception task between –1 and 1 seconds for the left hemisphere (1.5 < BF_10_ < 1.8) and between –3 to 4 seconds in the right hemisphere (3.9 < BF_10_ < 7). Given the hemodynamic delay, these responses in IFGs correspond to an anticipation period (before 0 s). The activation observed during the anticipation period in this experiment is likely attributable to the block design employed, where each block comprised trials of the same task and material. As in depth encoding of S1 is only needed in memory blocks, participants seemed to have adopted different attending strategies depending on the blocks. To avoid such block-related anticipatory effects, conditions were randomized within blocks in Experiment 2.

The analysis on HbR deconvoluted signal (supplementary Figure S3) mirrored HbO results in lateral frontal regions: we observed in the right dlPFC evidence for a stronger decrease of HbR for the memory task as compared to the perception task between 9 and 17 seconds after S1 onset (BF_10_ > 1) and in particular, 12 to 14 seconds after S1 onset (all BF_10_ > 13.4). We also observed a stronger decrease of HbR for the memory task compared to the perception task in the right IFG between –1 to 6 seconds (1 < BF_10_ < 114). Finally, no effect of task was observed for the SFG. Overall, these results are in accordance with previously reported findings that observed the involvement of lateral PFC areas in auditory memory processes, but not of medial frontal areas (Owen et al., 2005; Rottschy et al., 2012; Rovetti et al., 2021). They thus confirm that fNIRS is a suitable technique to explore the involvement of frontal areas in musical and verbal STM processes using a DMST paradigm.

## Experiment 2

### Methods

#### Participants

Twenty-four healthy adults were recruited for Experiment 2 (mean age = 28.9 years, sd = 9.3 years, min = 21 years, max = 66 years, 6 left handed, 16 females, mean education level = 15.44 years, mean musical education = 0.48 years, sd = 1 year). Participants underwent the same inclusion procedure as for Experiment 1. They all gave written informed consent to participate in the experiment (ethical authorization: CPP Ile de France VI, ID RDC 2018-A02670-55) and were given a compensation for their participation. The first session (∼ 1h30) was the same as for Experiment 1 and the second session during which participants underwent fNIRS recordings lasted around 1h30.

#### Stimuli and task design

In Experiment 2, only the STM task was used. There were three memory load (ML) levels that differed in sequence length: ML1, ML2, and ML3. For the musical material, the three MLs consisted in respectively, four-, five-, and six-item sequences; for the verbal material, they consisted in respectively six-, seven– and eight-item sequences. Within a trial, S1 and S2 always had the same number of items. Item duration, inter-stimulus interval, delay duration, response time-window and jittered inter-trial interval were as described for Experiment 1. Due to the ML manipulation, the durations of S1-delay-S2 were respectively 10600 ms, 11800 ms and 13000 ms for ML1, ML2, and ML3 for the musical material, and respectively 13000 ms, 14200 ms and 15400 ms for ML1, ML2, and ML3 for the verbal material.

Twelve S1 sequences were created for each ML level and for each material: six S1 sequences were used to create “same” trials, with S2 identical to S1. Six other S1 sequences were used to create “different” trials, with S2 sequences differing from S1. Only six items per material were available in order to maximize phonological discriminability and as the verbal stimuli are intended to be used in children with language disorders as well as in adults. Since there were sequences with more than six items, we allowed for item repetitions within a given sequence with the following constraints: the first item could not be repeated within a sequence, items could be repeated only with at least 2 items in-between, one item could not be repeated more than three times, and three– or four-item patterns could not be repeated. When S2 was different, two adjacent items were switched (instead of introducing a new item) thus systematically changing S2 contour for the musical material. Any item could be switched with the next one, except for the first item. When it was possible, there was an equiprobable number of sequences with each position of item switch (e.g., for musical ML1 “different” S2 sequences, there were three trials with a switch between the 2^nd^ and the 3^rd^ item and three trials with a switch between the 3^rd^ and the 4^th^ item). When it was not possible, the remaining number of trials was randomly assigned to an item position switch (e.g. for verbal ML1 “different” S2 sequences, there were five trials with an item-switch respectively between the 2^nd^, the 3^rd^, the 4^th^ and the 5^th^ item with the adjacent one and one randomly assigned trial with a switch between the 3^rd^ and the 4^th^ items). Additionally, twenty-four 26400– to 30400– ms randomly jittered silent trials were generated to intersperse within each testing block.

#### fNIRS data acquisition and montage

Data acquisition and fNIRS montage were the same as described in Experiment 1.

#### Procedure

Participants underwent six blocks of sixteen trials each. Each block could randomly contain any stimulation trial as there were only memory trials for Experiment 2 and we wanted to avoid condition-dependent anticipation effects (as observed in Experiment 1 with a block design). There could be no more than three consecutive “same” (or “different”) trials, no more than three consecutive trials of the same memory load, and no more than four consecutive trials of the same material. Four silent trials were displayed on the 1^st^, 6^th^, 11^th^ and 16^th^ trial position of each block. There was a six-trial training block at the beginning of the experiment, with one trial from each ML condition, half of them “same” and half of them “different”. Then, participants underwent the six consecutive test blocks. Each of the six blocks lasted around 6 minutes. The order of test blocks was counterbalanced across participants using a Latin square balanced for first-order carryover effects using the *crossdes* R package (Sailer, 2022).

#### Behavioral data analysis

Behavioral analysis was performed, as for Experiment 1, on d’, c, and correct RTs. For d’ and c, a Bayesian repeated-measure ANOVA was performed with Memory Load (three levels: ML1, ML2, ML3), Material (two levels: musical and verbal), their interaction as fixed factors and participants as random factor. Overall, four models were tested (memory load, material, memory load + material, memory load + material + memory load:material) and compared to the null model. Paired Bayesian t-tests were performed as post-hoc tests if the best model included the memory load effect or the interaction. Correct RTs were submitted to the same Bayesian ANOVA, with the addition of the type of trial (same/different) as factor. Additionally, one-sample Bayesian t-tests against 0 were performed on the criterion (c) for each memory load and material.

#### fNIRS data pre-processing and deconvolution

The raw data were preprocessed using the same pipeline as for Experiment 1. Data were then analyzed with the same FIR model as in Experiment 1. However, as S1 sequences had different durations according to memory load, we centered the deconvolution around the onset of the delay. Hence, 36 one-second boxcar regressors were fitted around delay onset (–9 seconds to 27 seconds). Overall, each recording channel of each participant was regressed using an Ordinary Least Squares (OLS) GLM with a design matrix including block regressors, 36 boxcar regressors around delay onsets for each memory load and material for the FIR deconvolution (216 regressors in total), and all short-channel signals (16 regressors).

#### fNIRS data analysis

The same ROIs as in Experiment 1 were used, with betas averaged across channels making up each ROI. These ROI-averaged betas were then tested across participants every second from –9 to 12 seconds around delay onset (i.e., until 6 seconds after the end of S2 for the highest ML) in each ROI with a repeated-measure Bayesian ANOVA including Memory Load (three levels: ML1, ML2, and ML3), Material (two levels: musical and verbal), and their interaction as fixed factors, with the exact same procedure as for the analysis of the behavioral data. Participants were added to all models as random factors with their random slopes for task and material. If the interaction effect was included in the best model, a Bayesian repeated-measure ANOVA with memory load as fixed factor and participants as random factor was performed separately for each material. Then for time samples for which the best model included the memory load effect for a given material, paired Bayesian t-tests were performed as post-hoc tests between each memory load levels (ML1 vs. ML2, ML1 vs. ML3, and ML2 vs. ML3).

As for Experiment 1, we report in the main text only results for HbO, and HbR results are available in supplementary materials (Figures S4) and commented in the discussion.

## Results

### Behavioral results

For d’ (Figure 4a), the best model explaining the data included both fixed factors (memory load and material, decisive evidence, BF_10_ = 6.1e+15) but not the interaction between both. The analysis of effects across matched models revealed decisive evidence for memory load (BF_inclusion_ = 2.83e+08) and material (BF_inclusion_ = 1.43e+10) factors. d’ was lower for the verbal material as compared to the musical material, and decreased as load increased (Figure 4a). Post-hoc Bayesian t-tests for the memory load factor revealed decisive evidence for a difference between ML1 and ML2 (BF_10_ = 3e+04) with lower d’ for ML2 as compared to ML1, decisive evidence for a difference between ML1 and ML3 (BF_10_ = 2.2e+05) with lower d’ for ML3 as compared to ML1, and weak evidence for a difference between ML2 and ML3 (BF_10_ = 2) with lower d’ for ML3 as compared to ML2.

**Figure 4:**
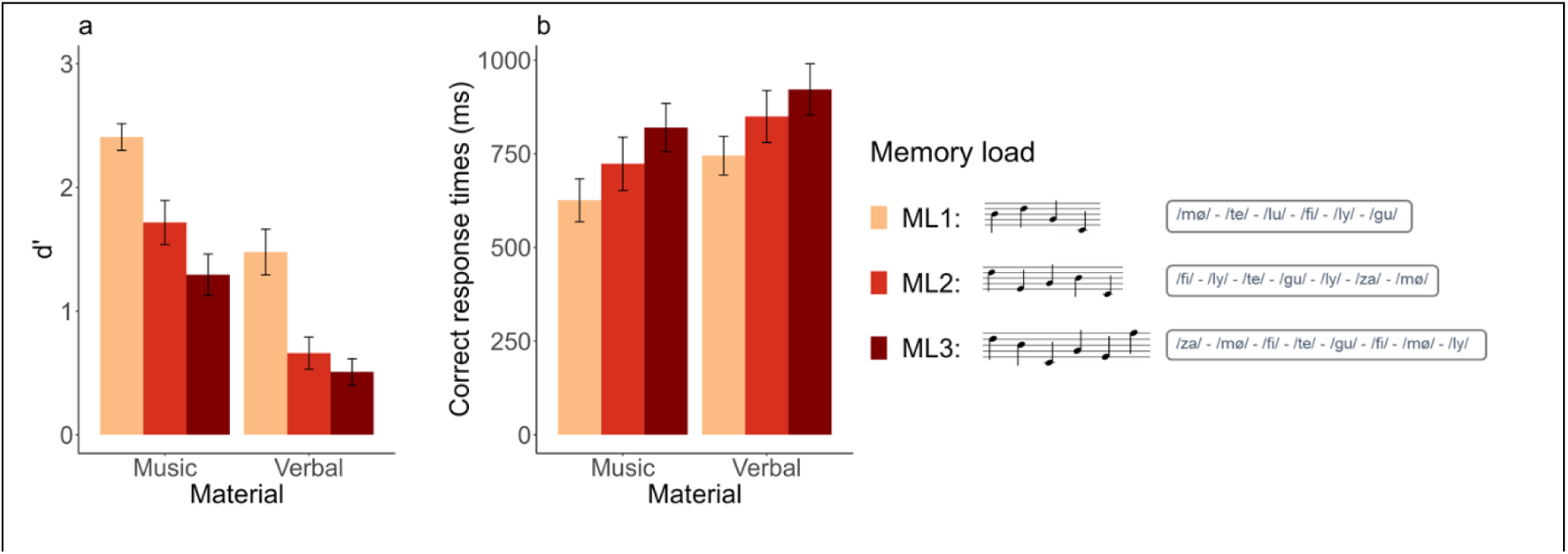
Average performance as a function of memory load (ML) (ML1 in orange/ML2 in red/ML3 in dark red) and material (musical/verbal). (a) Mean and standard error of participants’ sensitivity (d’). (b) Mean and standard error of averaged response times for correct trials (time in millisecond that participants spent after the end of S2 before giving a “same” or “different” answer). Examples of S1 sequences for each ML are shown in the legend.

For correct RTs (Figure 4b), the best model was, as for d’, the one including the fixed factors memory load and material (decisive evidence, BF10 = 1.4e+03). The analysis of effects across matched models revealed decisive evidence for the memory load factor (BF_inclusion_ = 2.3e+05) and strong evidence for the material factor (BF_inclusion_ = 65). RTs were longer for the verbal material as compared to the musical material, and increased with load (figure 4b). Post-hoc Bayesian t-tests for the memory load factor revealed positive evidence for a difference between ML1 and ML2 (BF_10_ = 8.8) with longer RTs for ML2 as compared to ML1, decisive evidence for a difference between ML1 and ML3 (BF_10_ = 1.3e+04) with longer RTs for ML3 as compared to ML1, and weak evidence for a difference between ML2 and ML3 (BF_10_ = 1) with slightly longer RTs for ML3 as compared to ML2.

For c, results are reported in supplementary Figure S4.

#### fNIRS results

HbO deconvoluted signals within the targeted ROIs are shown with corresponding statistics in Figure 5 (topographic representations are available in supplementary Figure S5 and HbR results in supplementary Figure S6). For completeness we report all effects below, but our interest was only in models including the memory load factor or its interaction with the material factor.

**Figure 5:**
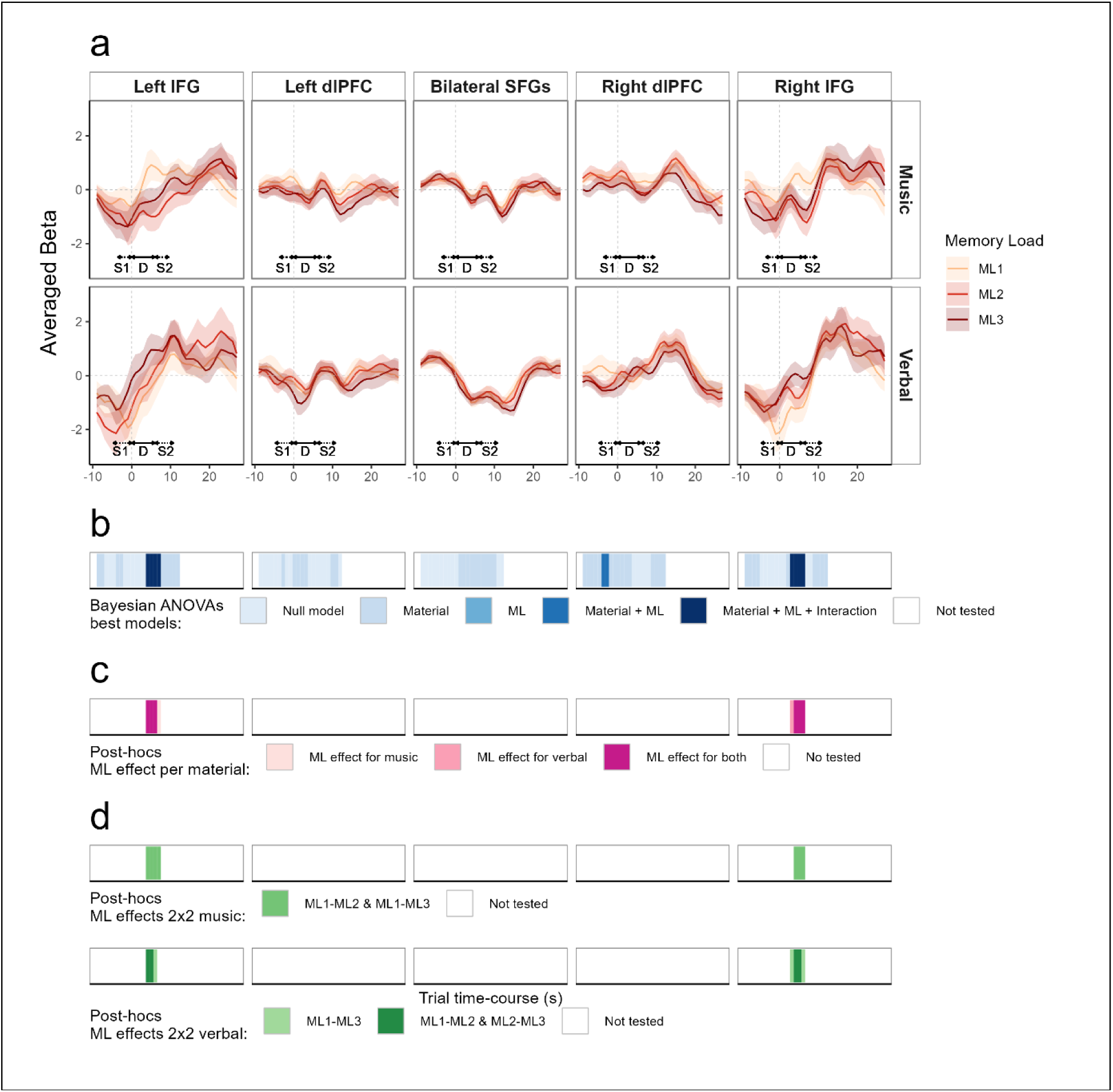
(a) Average beta (plain line) and standard error (shaded area) from the FIR deconvolution performed on HbO data for all participants (n=24), across five ROIs (left and right IFG, left and right dlPFC, bilateral SFGs), in a time window ranging from –9 to 27 seconds around delay onset (grey dotted vertical line), for the three memory load levels (ML1/ML2/ML3), for the musical material (top panel) and verbal material (bottom panel). For clarity purposes, S1, silent retention delay (D), and S2 durations are indicated with double-headed arrows, S1 and S2 arrows are dotted to indicate their variable duration according to the memory load. (b) Time-course representation of Bayesian analysis, each blue shade represents the best model as compared to the null model (BF_10_ > 1). No statistics were performed beyond 12 seconds after delay onset (end of S2 for the longest sequence + peak of the hemodynamic response, ∼ 6s). (c) Time-course representation of post-hoc Bayesian ANOVAs per material with memory load (ML) as factor performed only if the best model included the interaction. Purple shades indicate if there was evidence in favor of a memory load effect (BF_10_ > 1) for the musical material, the verbal material, or both. (d) Post-hoc Bayesian t-tests comparing memory load levels two by two for the musical material (top panel) and verbal material (bottom panel) performed only if the previous post-hoc analysis showed evidence for a memory load effect for the corresponding material.

For the left IFG, we found weak to strong evidence for the model with the material factor at time samples –9, –8, –4, and –3 seconds before delay onset (1.2 < BF_10_ < 18.6) with higher betas for the musical material as compared to the verbal one, and weak to strong evidence for the model with the material factor 8 to 12 seconds after delay onset (2 < BF_10_ < 39.4) with higher betas for the verbal material as compared to the musical one. Importantly, we found positive to strong evidence for the model including the interaction between material and memory load 4 to 7 seconds after delay onset (3.4 < BF_10_ < 46.4). In the musical material, post-hoc tests for the memory load effect revealed a positive to strong evidence for the memory load effect for the four tested time samples (10.3 < BF_10_ < 49.7). Pairwise post-hoc t-tests in the musical material revealed positive to strong evidence for a difference between ML1 and ML2 (4.4 < BF_10_ < 10.4) and weak to positive evidence for a difference between ML1 and ML3 (1.8 < BF_10_ < 4.9) for the four tested time samples. Unexpectedly, in the musical material, betas were higher for the ML1 condition as compared to the ML2 and ML3 conditions. In the verbal material, post-hoc tests for the memory load effect revealed weak to positive evidence for the memory load effect at time samples 4, 5, and 6 seconds (1.3 < BF_10_ < 5.3). Pairwise post-hoc tests in the verbal material revealed weak evidence for a difference between ML1 and ML3 (1.2 < BF_10_ < 2.4) for the three tested time samples and weak evidence for a difference between ML2 and ML3 (1.2 < BF_10_ < 1.5) for time samples 4 and 5. As expected, for the verbal material, betas were higher for the ML3 condition as compared to the ML1 and ML2 conditions. In other words, higher activations were found for the ML1 condition as compared to ML2 and ML3 in the musical material, and higher activations were found for the ML3 condition as compared to ML2 and ML1 in the verbal material.

For the right IFG, we found weak to positive evidence for the model including the material factor –9 to –6 seconds and at 2 seconds around delay onset (1.1 < BF_10_ < 10) with higher betas for the musical material as compared to the verbal one. Importantly, we found weak to strong evidence for the model including the interaction 3 to 6 seconds after delay onset (1 < BF_10_ < 19.8). Post-hoc tests for the memory load effect in the musical material revealed a weak to positive evidence for the memory load effect for time samples 4, 5, and 6 seconds (2.1 < BF_10_ < 7.7). Pairwise post-hoc bayesian t-tests in the musical material and for these three time samples revealed weak to positive evidence for a difference between ML1 and ML2 (1.2 < BF_10_ < 3.5) and weak to positive evidence for a difference between ML1 and ML3 (2.8 < BF_10_ < 5). As for the left IFG for the musical material, betas were higher for the ML1 condition as compared to the ML2 and ML3 conditions. Post-hoc tests for the memory load effect in the verbal material revealed weak to strong evidence for the memory load effect for the four tested time samples (1.5 < BF_10_ < 15.7). Pairwise post-hoc t-tests in the verbal material revealed weak to positive evidence for a difference between ML1 and ML3 (1.6 < BF_10_ < 7.8) for the four tested time samples, and positive evidence for a difference between ML2 and ML3 (3.9 < BF_10_ < 4.9) at time samples 4 and 5 seconds. As expected, for the verbal material, betas were higher for the ML3 condition as compared to the ML1 and ML2 conditions.

For the left dlPFC, we found weak to strong evidence for the model including the material factor at –3 seconds and from 0 to 3 seconds around delay onset (1 < BF_10_ < 11.2) with higher betas for the musical material as compared to the verbal one and weak evidence for the model with the material factor 10 and 11 seconds after delay onset (1.2 < BF_10_ < 1.4) with higher betas for the verbal material as compared to the musical one.

For the right dlPFC, we found weak to strong evidence for the model including the material factor at –9 to –5 seconds and –2 to 3 seconds around delay onset (1.2 < BF_10_ < 19.1) with higher betas for the musical material as compared to the verbal one. We also found weak to strong evidence for the model with the material factor 9 to 12 seconds after delay onset (1.3 < BF_10_ < 15.2) with higher betas for the verbal material as compared to the musical one. We found strong evidence for the model including material and memory load as factors –4 and

-3 seconds before delay onset (20.4 < BF_10_ < 44.2) with higher betas for the musical material as compared to the verbal one. Post-hoc tests averaged over materials for the memory load effect (not shown in Figure 7) revealed weak evidence for a difference between ML1 and ML2 (1.4 < BF_10_ < 1.9) and positive to strong evidence for a difference between ML1 and ML3 (9.8 < BF_10_ < 10.1), with higher betas for ML1 as compared to ML2 and ML3.

For the SFG, we found weak to strong evidence for the model including the material factor from 1 to 10 seconds after delay onset (1.1 < BF_10_ < 22.8) with higher betas for the musical material as compared to the verbal material.

## Discussion

The aim of Experiment 2 was to manipulate memory load for musical and verbal STM in order to identify the lateral frontal regions showing parametric involvement with memory load and to explore whether their involvement differs for the two materials.

Behavioral data showed a parametric decrease of performance and an increase of RTs for higher memory loads with strong to decisive evidence for differences between ML1 and the other two memory loads (Figure 4), as previously observed in studies manipulating memory load with DMST (Grimault et al., 2014; Schulze et al., 2012; Schulze & Tillmann, 2013). Analysis of the criterion (c) revealed weak evidence for a liberal bias for low memory loads for the musical material. The performance was overall lower and RTs longer for the verbal material as a consequence of (1) increasing sequence length by two items in the verbal material compared to the musical material and (2) switching from a change of item to a change of order when S2 sequences were different from S1. In future studies using these stimuli, similar performance between musical and verbal material should be reached by increasing sequence length for the verbal material by one item compared to the musical material. Nevertheless, we managed to manipulate memory load in a similar fashion for both materials, as revealed by the absence of interaction between material and memory load for both performance and RTs. These results provide evidence that the DMST paradigm is highly effective in manipulating memory load, in line with previous behavioral studies with auditory material (Schulze et al., 2012; Schulze & Tillmann, 2013) and the few neuroimaging studies employing DMST for this purpose in the visual (Habeck et al., 2005; Kaiser et al., 2010; Robitaille et al., 2010) and auditory (Grimault et al., 2014) modalities.

As S1 duration was different for each memory load (different sequence lengths), rendering hemodynamic time-course comparison in the response to S1 difficult, we analyzed the hemodynamic responses from the start of the retention delay. Note that material effects in the time ranges corresponding to S1 encoding or S2 processing are likely reflecting the differences in sequence lengths for the two materials. Hence, the interpretation of the data focuses on the delay period. The topographical representation of HbO deconvoluted signal (supplementary Figure S5) showed a parametric modulation of activity in lateral frontal channels for both materials after the delay onset with higher activation for lower memory loads for the musical material and, conversely, higher activation for higher memory loads for the verbal material. In the ROI analysis (Figure 5), the parametric effects in the IFGs were present 3 to 6 seconds after delay onset but not later, confirming that the parametric effect specifically concerned processes at the beginning of the silent retention delay. For the verbal material, we observed the expected activation increase with higher memory loads but intriguingly, for the musical material, the effect of memory load was not the expected one, with higher activation for ML1 as compared to ML2 and ML3. We discuss these results, along with those from Experiment 1, in the following general discussion.

Channels placed along the medial part of the PFC (SFGs) were not sensitive to memory load, as expected. They showed a global deactivation after delay onset regardless of the memory load, but more strongly for verbal stimuli. In the ROI analysis (Figure 5), we observed that the dlPFC and SFG did not show any memory load effect, and that the parametric memory load effects only involved the bilateral IFGs. Furthermore, in the SFG, the material effect revealed greater deactivation for the verbal material, likely reflecting the increased difficulty of the task, as revealed by the lower performance for the verbal material than for the musical material.

## General Discussion

The aims of the present study were three-fold. (1) We aimed at validating the application of fNIRS in investigating frontal activations during auditory DMST. In Experiment 1, the IFG and dlPFC exhibited greater activation for the memory task as compared to the perception task for both materials while the SFG did not exhibit any task effect. (2) Our second goal was to identify the specific frontal brain regions where activity demonstrates parametric variation with memory load. In Experiment 2, only the IFG showed parametric activations with memory load while the dlPFC and the SFG did not respond to memory load manipulation. (3) We wanted to determine whether these regions exhibit similarities when processing musical and verbal materials. Both materials showed similar patterns in Experiment 1 (memory > perception in the IFG and the dlPFC) and memory load processing involved the same region (IFG) in Experiment 2. However, different profiles were observed between materials with increasing activations for increasing memory loads in the verbal material and decreasing activation for increasing memory loads in the musical material.

### Involvement of frontal regions in auditory STM

Previous research has consistently indicated the involvement of lateral prefrontal regions during maintenance of musical and verbal material in auditory STM, specifically the IFG and dlPFC. None of the previous studies has reported the involvement of medial prefrontal regions (Grimault et al., 2014; Rottschy et al., 2012). In the current study, both experiments provide converging evidence with these observations. In particular, Experiment 1 provided (1) evidence for the involvement of bilateral IFGs during the anticipation period (–3 to 2 seconds around S1 onset), only for the memory task. These results are an addition to previous fMRI studies that did not investigate anticipation effects. (2) During the maintenance of information (i.e., the silent retention delay), we found that the dlPFC showed higher activation for the memory task as compared to the perception task, while this pattern was not observed in the IFG. The findings in previous studies present inconsistencies regarding the specific involvement of the dlPFC or IFG in the maintenance of musical and verbal STM. While Koelsch et al. (2009) observed activations in the IFG, but not the dlPFC, during the maintenance phase for musical and verbal information, Schulze et al. (2011) and Albouy et al. (2019) reported involvement of both regions during the maintenance period. Given that our study is the first to directly compare materials using a neuroimaging technique that differs from fMRI, combined with the scarcity of existing research in this area, further studies are necessary to draw conclusive insights regarding whether the IFG and/or the dlPFC is preferentially involved in maintenance in auditory STM tasks.

In Experiment 1, our findings revealed, in line with previous fMRI studies, that when comparing memory processes to perception processes in control participants, no difference between musical and verbal material was observed. In addition, we did not observe differences in the medial frontal regions activity for memory processes compared to perception processes. Overall, by demonstrating a differential activation patterns between memory and perception tasks in lateral PFC, our study contributes to the growing body of evidence linking lateral prefrontal regions to the encoding and maintenance of auditory information in STM, a link that was further explored in Experiment 2 by manipulating memory load.

### Memory load manipulation for musical and verbal STM

In Experiment 2, our results demonstrated that, as the verbal memory load increased, there was a corresponding increase in activation in bilateral IFGs during the maintenance of information. In contrast, the verbal memory load did not impact bilateral dlPFC and SFG activations. Previous studies in fMRI report a linear increase in activity with visual verbal memory load using n-back tasks (assessing WM) in the left IFG and in bilateral dlPFC (Braver et al., 1997) and in the left IFG and right dlPFC (Cohen et al., 1997). fNIRS studies have also yielded similar results for visual-verbal memory load using n-back tasks (Fishburn et al., 2014; Khaksari et al., 2019). Fishburn et al. (2014) showed an increase in dlPFC with load but not in the IFG, medial PFC, or parietal areas. Khaksari et al (2019) found an increase of activity in bilateral dlPFC memory load (they did not record bilateral IFGs). Interestingly, they also found a parametric increase in medial PFC. However, these results can be mitigated as most of the fNIRS studies analyzed HbO and HbR data, whereas in this study, the authors only analyzed total hemoglobin concentration changes, making the comparison difficult. In the auditory modality, fNIRS studies have found an increase of activity with verbal WM load in bilateral IFGs and dlPFCs (without differentiating them in the fNIRS montage, Rovetti et al., 2021) and in bilateral orbital PFC and IFG, but without recording bilateral dlPFCs (Tseng et al., 2018).

Overall, for verbal material, WM studies consistently report a linear bilateral increase in lateral prefrontal regions with memory load, but most of the time not in medial frontal regions. The relative involvement of the dlPFC and the IFG varies across studies. One potential explanation for the present findings implicating mostly the IFG is that our DMST paradigm targeted STM rather than WM processes. With fMRI, using forward (STM) and backward (WM) digit span tasks, the orbital part of the IFG showed significant activation for both tasks while the dlPFC showed significant activation only for the WM task. With fNIRS, the contrast between forward and backward digit span tasks shows that only the dlPFC is involved in WM processes (Tian et al., 2014). The relative involvement of IFG, dlPFC, and SFG might thus depend on task requirements, neuroimaging technique, modality, and analysis pipelines.

For musical STM, our study revealed that during the maintenance of information, activity varied parametrically with load in bilateral IFGs but not in bilateral dlPFCs nor SFGs. However, in contrast to the parametric variation for verbal material, we found a parametric decrease of activation with the increase of memory load and more specifically, higher activations for the 1^st^ level of memory load (4-item sequences) as compared to levels 2 and 3 (5– and 6-item sequences). These results are not in line with the only study that investigated memory load effects for musical material using MEG: Grimault et al. (2014) reported an increase of activity with memory load in bilateral IFGs, dlPFCs and temporal regions. The main difference between this latter study and the present one, beyond the use of different neuroimaging techniques, is that they used 4-item sequences as their maximum memory load, when in the present study a 4-item sequence corresponded to the minimum memory load, 6-item sequences being the maximum one. Furthermore, their silent retention interval lasted 2 seconds when ours lasted 6 seconds. Therefore, the task used in the present study was far more challenging than the task used in Grimault et al. (2014). While studies employing verbal n-back tasks have reported deactivation in lateral prefrontal regions when memory loads exceed participants’ WM capacity (Nyberg et al., 2009), we can exclude this explanation for the present study. Indeed, our behavioral results demonstrate that participants were able to perform the musical task even at the highest memory load, and we observed a parametric increase of IFG activity with load in the more difficult verbal task. A possible explanation for these findings could be that when processing musical sequences in the lower memory load condition, participants employ the same strategy as for the verbal material and thus engage the same frontal network, as evidenced by the increased activity in the IFGs in the ML1 condition. However, for higher memory loads, participants might change strategy. It has been shown that for the encoding of verbal information, when the acoustic context facilitates chunking strategies, activity decreases in lateral frontal regions (Ferreri et al., 2015). Such activity decreases in lateral PFC are also observed when participants are given specific instructions to use a chunking strategy (Matsui et al., 2007). For musical material, participants might have relied on different strategies to handle higher memory loads. A previous study comparing musicians and non-musicians has evidenced the existence of different (and more or less efficient) strategies for musical STM, notably using or not contour information (Talamini et al., 2021). Some of these alternative strategies (chunking, contour-based, etc.) could involve different brain regions than the ones recorded here. Interestingly, when testing the impact of memory load for musical material with MEG, significant clusters of activation were observed not only in frontal areas, but also in temporal and parietal areas (Grimault et al., 2014). Future studies using musical material and a DMST should gather data about the strategy employed by participants to perform the task and record activity in temporal and parietal areas. Moreover, these maintenance strategies could be studied for other auditory material, such as timbre. Indeed, it has been suggested that strategies used to maintain for timbre information in WM differs from tonal and verbal material and that they would rely on sensory imagery rather than internal rehearsal or moto-related processes (Schulze & Tillmann, 2013).

Previous neuroimaging studies comparing musical and verbal STM consistently found overlapping regions involved for both materials, except in impaired or expert populations (Caclin & Tillmann, 2018). Our results support the observation that the same brain regions are recruited for musical and verbal STM, but by manipulating memory load, we found that their involvement differed between the two materials. These results yield great promise for the identification of specific neurophysiological markers of auditory STM for specific materials. Here, by increasing the length of the verbal sequences by two items compared to the musical sequences, we managed to have a similar influence of increases in memory loads on behavior, but with an overall decrease of performance for verbal sequences compared to musical ones. To go further, future studies should equalize performance levels between the two materials, along with recording other regions that have been shown to be involved in memory processes (e.g. temporal and parietal regions). One potential limitation of fNIRS is its limited ability to record deeper brain regions, such as the primary auditory cortex and surrounding areas, which could also be involved in musical memory. While some studies have reported success in detecting auditory cortex activity using fNIRS (Plichta et al., 2011; Santosa et al., 2014), it remains a methodological challenge. Additionally, it would be valuable to investigate the activity of parietal regions, such as the superior and inferior parietal lobule, which activity has been shown to be modulated by memory load for musical and verbal material (Grimault et al., 2014; Rovetti et al., 2021).

### Perspectives for studies in children with learning disorders

fNIRS is a promising tool for investigating auditory cognition in children, with typical development and with learning disorders, such as dyslexia and DLD, thanks to its portability, non-invasiveness, and silent nature compared to fMRI. As we have demonstrated in the present study, fNIRS is effective in measuring neurophysiological markers of auditory STM and can differentiate the effects of memory load between different materials. To our knowledge, only two studies have investigated neural activity in children with developmental language disorder (DLD) with fNIRS so far. Decreased HbO activation in bilateral IFG and parietal regions can be observed in children with DLD during a language comprehension task compared to typically developing (TD) children (Fu et al., 2016) while reduced engagement of the left dlPFC and bilateral IPLs can be found in DLD children during a verbal WM task, compared to TD children. Overall, these previous studies and the current findings highlight the potential of fNIRS as a suitable neuroimaging technique to explore functional differences between children with DLD and TD children, and suggest that variations of cerebral activity in response to memory load manipulation could be an adequate neurophysiological marker. Future studies could investigate further the engagement of lateral frontal regions during auditory STM for musical and verbal material in children with DLD as deficits in STM for both materials have been observed (Couvignou et al., 2023; Couvignou & Kolinsky, 2021; Forgeard et al., 2008; Ziegler et al., 2012). In the long term, a better understanding of the neural mechanisms underlying auditory STM in DLD and their link to the processes reported to be impaired in this population (e.g., syntax processing, phoneme discrimination, etc.) could lead to the development of more effective and targeted interventions for this population.

## Data and code availability statement

The data and the code that support the findings of this study are available on request from the corresponding author.

## CRediT authorship contribution statement

**Jérémie Ginzburg**: Conceptualization, Investigation, Methodology, Visualization, Resources, Software, Writing – original draft, Writing – review & editing. **Anne Cheylus**: Methodology, Software, Writing – review & editing. **Elise Collard**: Investigation, Writing – review & editing. **Laura Ferreri**: Conceptualization, Writing – review & editing, Funding acquisition. **Barbara Tillmann**: Conceptualization, Resources, Writing – review & editing, Funding acquisition. **Annie Moulin**: Conceptualization, Resources, Writing – review & editing, Funding acquisition. **Anne caclin**: Conceptualization, Supervision, Methodology, Resources, Software, Writing – review & editing, Funding acquisition.

## Acknowledgments

We thank Lilou Spinosi, Solène Houdeline, Laurine Milon, and Julie Robin for their help with data recording, as well as Federico Curzel for helpful discussions and Lesly Fornoni for her help with ethics and participants recruitment.

## Supplemental material

### fNIRS montage: channel specificity

**Table S1:** Each of the 22 recording channels is composed of a source and a detector (located in standard 10-20 positions). We report in the table the two cortical structures for which each channel has the highest specificity (in percent) according to the fOLD software (Zimeo Morais et al., 2018). SFG: Superior Frontal Gyrus, MFG: Middle Frontal Gyrus, IFG: Inferior Frontal Gyrus. The channels forming the IFG, the dlPFC (corresponding to MFG), and the SFG ROIs are highlighted with light blue, light gold, and grey colors respectively. Channels unused in the ROI analysis are not highlighted.

| Hemisphere | Source | Detector | Cortical structure | Specificity (%) |
| --- | --- | --- | --- | --- |
| Left | AF7 | F5 | IFG (p. Triangularis) | 54.20 |
|  |  |  | MFG | 25.02 |
|  | F7 | F5 | IFG (p. Triangularis) | 82.28 |
|  |  |  | IFG (p. Orbitalis) | 11.66 |
|  | FC5 | F5 | IFG (p. Triangularis) | 68.70 |
|  |  |  | IFG (p. Opercularis) | 19.56 |
|  | AF3 | F5 | MFG | 66.90 |
|  |  |  | IFG (p. Triangularis) | 22.08 |
|  | F3 | F1 | MFG | 68.06 |
|  |  |  | SFG | 30.84 |
|  | F3 | F5 | MFG | 60.23 |
|  |  |  | IFG (p. Triangularis) | 38.27 |
|  | F3 | FC3 | MFG | 81.08 |
|  |  |  | PG | 9.36 |
|  | AF3 | AFz | SFG | 46.49 |
|  |  |  | SFG, medial | 37.44 |
|  | Fz | F1 | SFG, medial | 40.89 |
|  |  |  | SFG | 39.90 |
|  | FC5 | FC3 | Precentral Gyrus | 46.40 |
|  |  |  | IFG (p. Opercularis) | 18.71 |
| Right | AF8 | F6 | IFG (p. Triangularis) | 38.12 |
|  |  |  | MFG | 37.98 |
|  | F8 | F6 | IFG (p. Triangularis) | 73.88 |
|  |  |  | IFG (p. Orbitalis) | 18.76 |
|  | FC6 | F6 | IFG (p. Triangularis) | 59.66 |
|  |  |  | IFG (p. Opercularis) | 25.22 |
|  | AF4 | F6 | MFG | 69.69 |
|  |  |  | SFG | 14.98 |
|  | F4 | F2 | MFG | 65.21 |
|  |  |  | SFG | 31.75 |
|  | F4 | F6 | MFG | 59.39 |
|  |  |  | IFG (p. Triangularis) | 39.41 |
|  | F4 | FC4 | MFG | 70.58 |
|  |  |  | IFG (p. Opercularis) | 10.29 |
|  | AF4 | AFz | SFG | 42.17 |
|  |  |  | SFG, medial | 36.66 |
|  | Fz | F2 | SFG, medial | 40.68 |
|  |  |  | SFG | 35.24 |
|  | FC6 | FC4 | Precentral Gyrus | 45.52 |
|  |  |  | IFG (p. Opercularis) | 30.53 |
| Interhemispheric | Fz | AFz | SFG, medial | 43.16 |
| sulcus | Fz | FCz | SFG, medial | 23.98 |

### Experiment 1

#### Behavioral results: criterion

**Figure S1:**
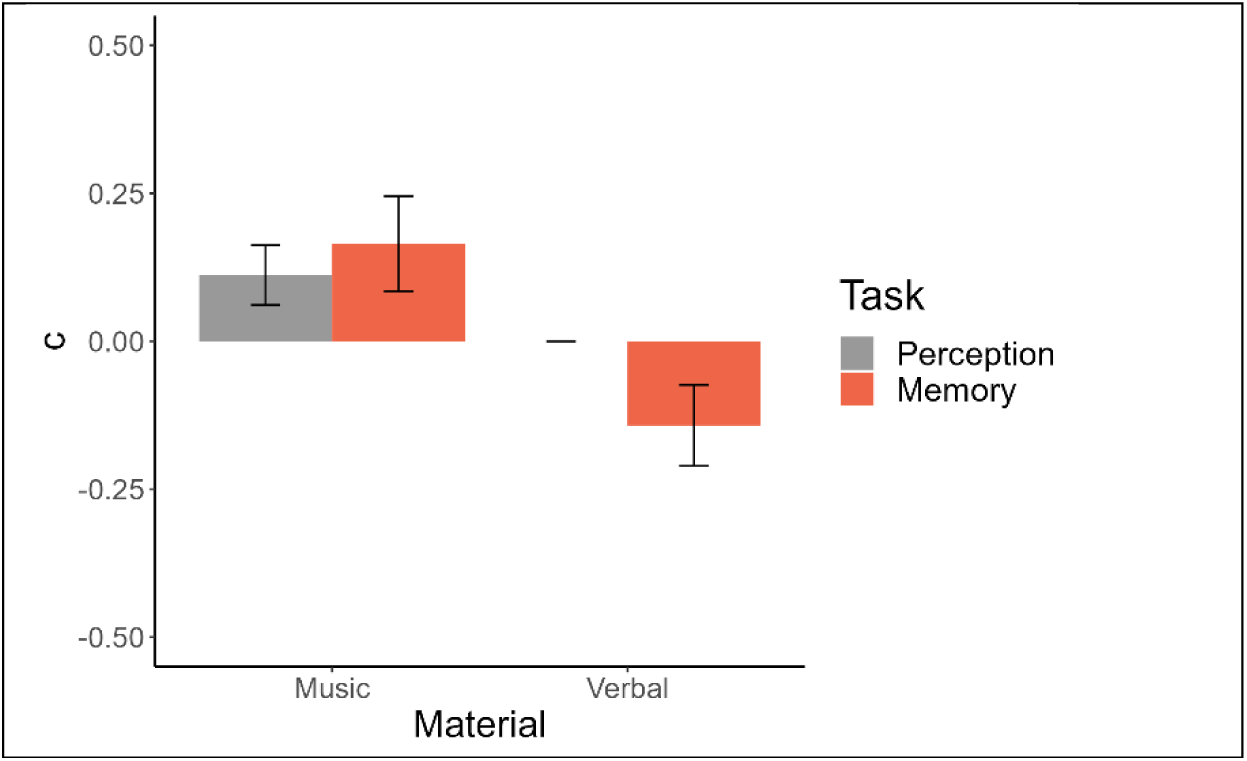
Mean and standard error of criterion (c) as a function of the task (perception in grey/memory in orange) and material (music/verbal).

For the criterion (c), the best model explaining the data included only the material factor (strong evidence, BF_10_ = 89.2) with a more conservative criterion for the musical material than for the verbal material. One-sample Bayesian t-tests revealed weak evidence for a difference compared to 0 for the perception and memory tasks in the musical material and for the memory task in the verbal material (1.3 < BF_10_ < 1.7). Note that for the perception task in the verbal material, all participants displayed a criterion of 0 (no bias) because they performed correctly for all materials.

#### Topographic representation of HbO results

**Figure S2:**
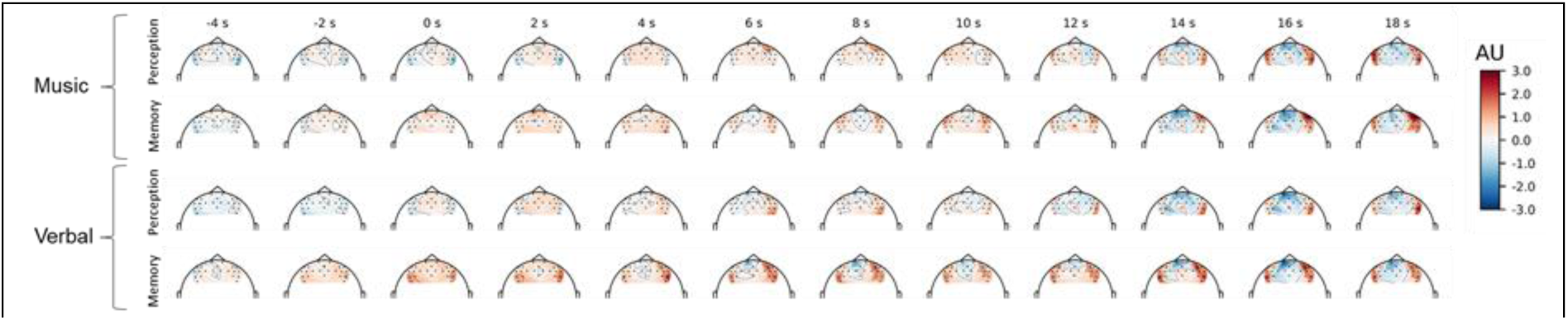
time-course of the topographic representation of the deconvoluted HbO fNIRS signal. Averaged beta across participants are represented every two seconds from –4 to 18 seconds around S1 onset for each channel, each task (perception/memory) and each material (music/verbal). AU: arbitrary units.

#### HbR results within targeted ROIs

HbR results within the targeted ROIs are summarized in Figure S2.

For the left IFG, we found weak evidence at –5 and –4 seconds before S1 onset for the model including the material effect (1 < BF_10_ < 1.6) higher betas for the verbal material as compared to the musical material.

For the right IFG, we found weak evidence –5 seconds before S1 onset for the model including the material effect (BF_10_ = 2.3) with higher betas for the verbal material as compared to the musical material. We found weak to decisive evidence –1 to 6 seconds around S1 onset for the model including the task effect (1.4 < BF_10_ < 160.1) with lower betas for the memory task as compared to the perception task.

For the left dlPFC, we found weak to strong evidence –5 to 3 seconds around S1 onset for the model including the task effect (1.1 < BF_10_ < 28.2) with lower betas for the memory task as compared to the perception task.

For the right dlPFC, we found weak to strong evidence –2 to 2 seconds and 9 to 17 seconds around S1 onset for the model including the task effect (1.3 < BF_10_ < 13.7) with lower betas for the memory task as compared to the perception task.

For the SFG, we found weak to positive evidence –2 to 1 seconds around S1 onset for the model including the task effect (1.6 < BF_10_ < 3.3) with lower betas for the memory task as compared to the perception task.

**Figure S3:**
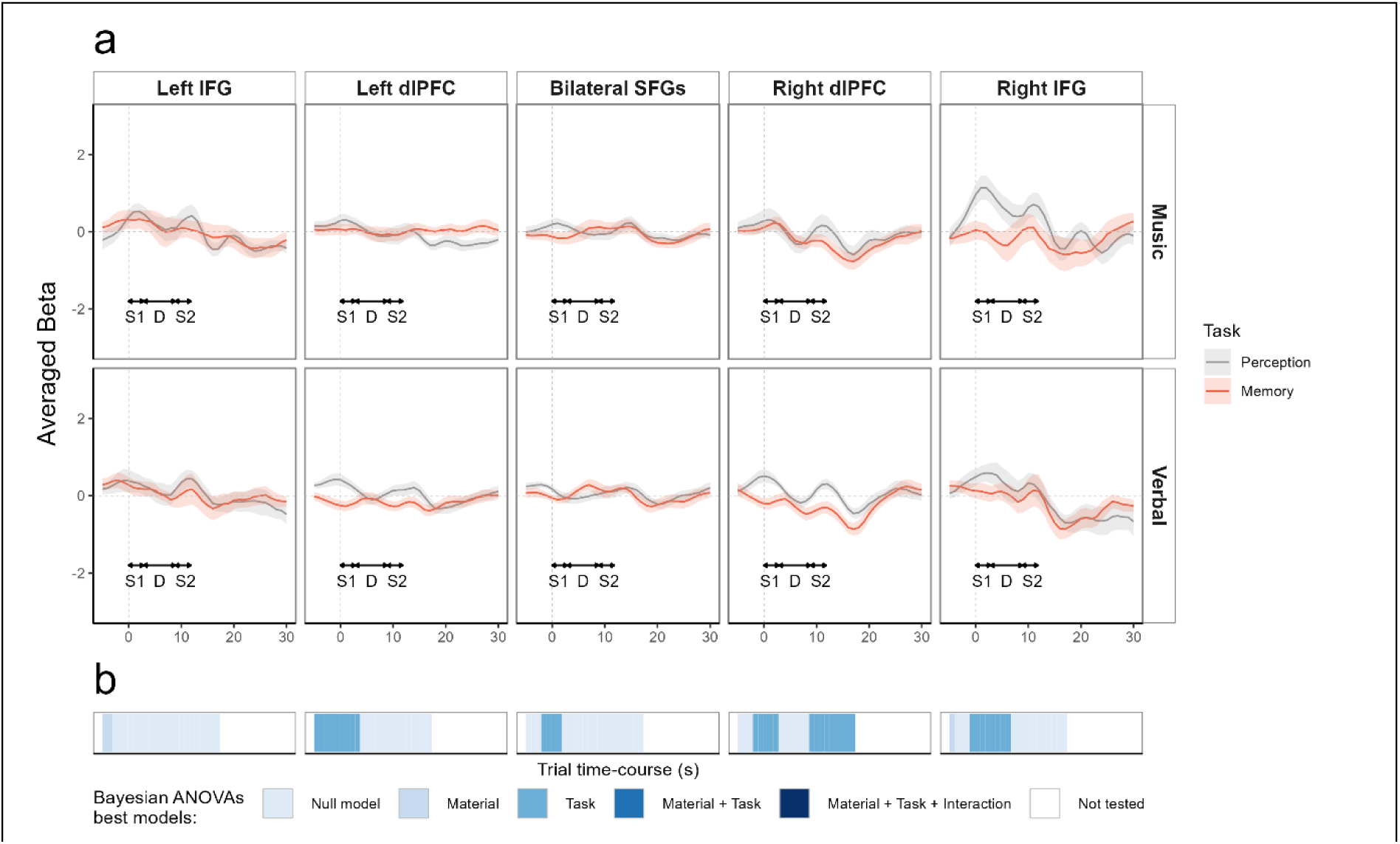
(a) Average beta (plain line) and standard error (shaded area) from the FIR deconvolution performed on HbR data for all participants (n=16), across five ROIs (left and right IFG, left and right dlPFC, bilateral SFGs), in a time window ranging from –5 to 30 seconds around S1 onset (grey dotted vertical line), for the perception task (grey) and the memory task (red). Top panel: musical material; bottom panel: verbal material. For clarity purposes, S1, delay (D), and S2 durations are indicated with double-headed arrows. (b) Time-course representation of Bayesian analysis, each blue shade represents the best model as compared to the null model (BF_10_ > 1). No statistics were performed after 18 seconds (end of S2 + peak of the hemodynamic response, ∼ 6s), because the deconvoluted signal would then include motor responses which we did not intend to analyze.

### Experiment 2

#### Behavioral results: criterion

**Figure S4:**
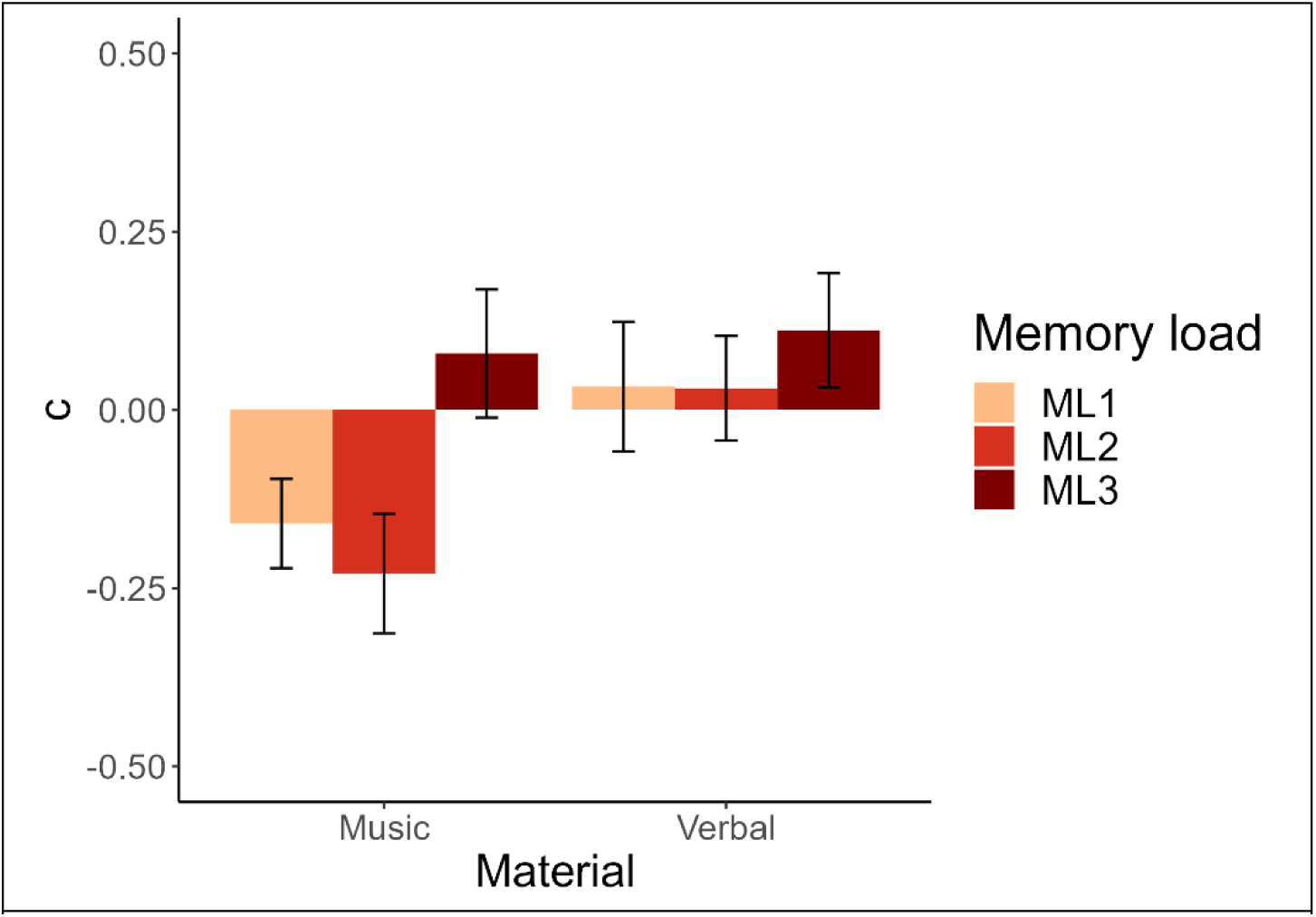
Mean and standard error of criterion (c) as a function of memory load (ML1 in orange/ML2 in red/ML3 in dark red) and material (music/verbal).

For the criterion (c), the best model explaining the data included both the condition and the material factor (positive evidence, BF_10_ = 6.3). The analysis of effects across matched models revealed weak evidence for the condition effect (BF_inclusion_ = 1.72) and positive evidence for the material effect (BF_inclusion_ = 3.5). For the material effect, participants displayed a more conservative criterion for the verbal material as compared to the musical material. Post-hoc Bayesian t-tests for the memory load factor averaged across materials revealed positive evidence for the null model between ML1 and ML2 (BF_10_ = 0.24), weak evidence for a difference between ML1 and ML3 (BF_10_ = 2.5) with a more conservative criterion for ML3 than for ML1, and positive evidence for a difference between ML2 and ML3 (BF_10_ = 8) with a more conservative criterion for ML3 than for ML2. One-sample Bayesian t-tests revealed weak evidence for a difference compared to 0 revealed positive evidence for a difference against for ML1 and ML2 conditions in the musical material (3 < BF_10_ < 4.2). For all other comparisons, weak to positive evidence for the null model (no difference compared to 0) was found (0.2 < BF_10_ < 0.5).

#### Topographic representation of HbO results

**Figure S5:**
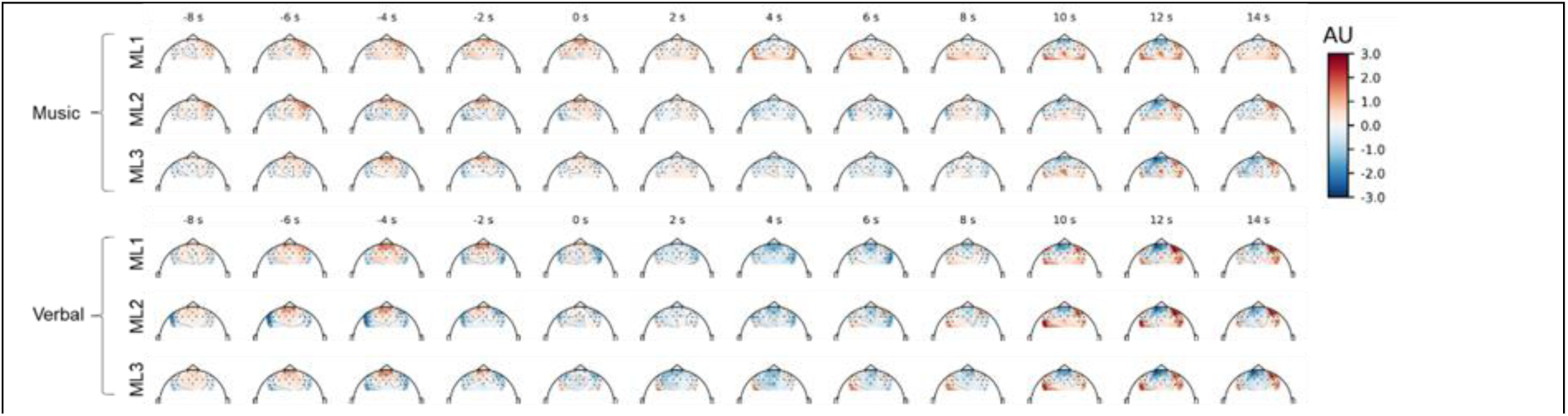
Time-course of the topographic representation of the HbO deconvoluted fNIRS signal. Averaged beta across participants are represented every two seconds from –8 to 14 seconds around delay onset (time sample 0) for each channel, each memory load (ML1/ML2/ML3), and each material (music/verbal). AU: arbitrary units.

#### HbR results within targeted ROIs

For the left IFG we found weak evidence at –8 seconds before delay onset for the model including the material effect (BF_10_ = 1.3) with lower betas for the musical material as compared to the verbal material. We found also weak to positive evidence at 3 and 4 seconds and 10 to 12 seconds after delay onset for the model including the material effect (1.1 < BF_10_ < 4.7) with lower betas for the verbal material as compared to the musical material.

For the right IFG we found weak evidence at –9 and –8 seconds before delay onset for the model including the material effect (1 < BF_10_ < 1.4) with lower betas for the verbal material as compared to the musical material. We found weak to strong evidence –2 to 4 seconds and 7 to 12 seconds around delay onset for the model including the memory load effect (1 < BF_10_ < 28.6). Post-hoc tests averaged over materials for the memory load effect (not shown in Figure S6) revealed weak to positive evidence 0 to 4 seconds after delay onset1 for a difference between ML1 and ML2 (1.4 < BF_10_ < 7.7) with lower betas for the ML2 condition as compared to the ML1 one. We found weak evidence 7 to 12 seconds after delay onset for a difference between ML1 and ML3 (1.5 < BF_10_ < 2.8) with lower betas for the ML3 condition as compared to ML1 condition. Finally, we found weak to positive evidence for all tested time samples (–2 to 4 and 7 to 12 seconds) for a difference between ML2 and ML3 (2.1 < BF_10_ < 5.9) with lower betas for the ML2 condition as compared to the ML3 condition.

For the left dlPFC, we found weak to positive evidence –2 to 2 seconds around delay onset for the model including the material effect (1.5 < BF_10_ < 8) with lower betas for the musical material as compared to the verbal material.

For the right dlPFC, we found weak to strong evidence at –9 seconds, –4 to 0 seconds and 10 to 12 seconds around delay onset for the model including the material effect (1 < BF_10_ < 49.6) with lower betas for the musical material –9 seconds and –4 to 0 seconds as compared to the verbal material and lower betas for the verbal material 10 to 12 seconds after delay onset as compared to the musical material. We found weak evidence –8 and –7 seconds before delay onset for the model including the interaction between material and memory load (1.5 < BF_10_ < 1.7). Post-hoc tests for the memory load effect in the musical material revealed a positive to strong evidence for the memory load effect for the two tested time samples (15 < BF_10_ < 24). Pairwise post-hoc t-tests in the musical material revealed strong evidence for a difference between ML2 and ML3 (19.3 < BF_10_ < 22.3) with lower betas for the ML2 condition as compared to the ML3 condition. Finally, we found weak to positive evidence 1 to 4 seconds after delay onset for the model including the memory load effect (1.7 < BF_10_ < 9.2). Post-hoc tests averaged over materials for the memory load effect (not shown in Figure S6) revealed positive evidence in all tested time samples for a difference between ML1 and ML2 with lower betas for ML2 as compared to the ML1 condition. There was also weak evidence at 1 and 2 seconds for a difference between ML2 and ML3 (1.1 < BF_10_ < 1.3) with lower betas for the ML2 condition as compared to the ML3 condition.

For the SFG, we found weak to strong evidence –1 to 10 seconds around delay onset for the model including the material effect (1 < BF_10_ < 74.9) with lower betas for the musical material as compared to the verbal material.

**Figure S6:**
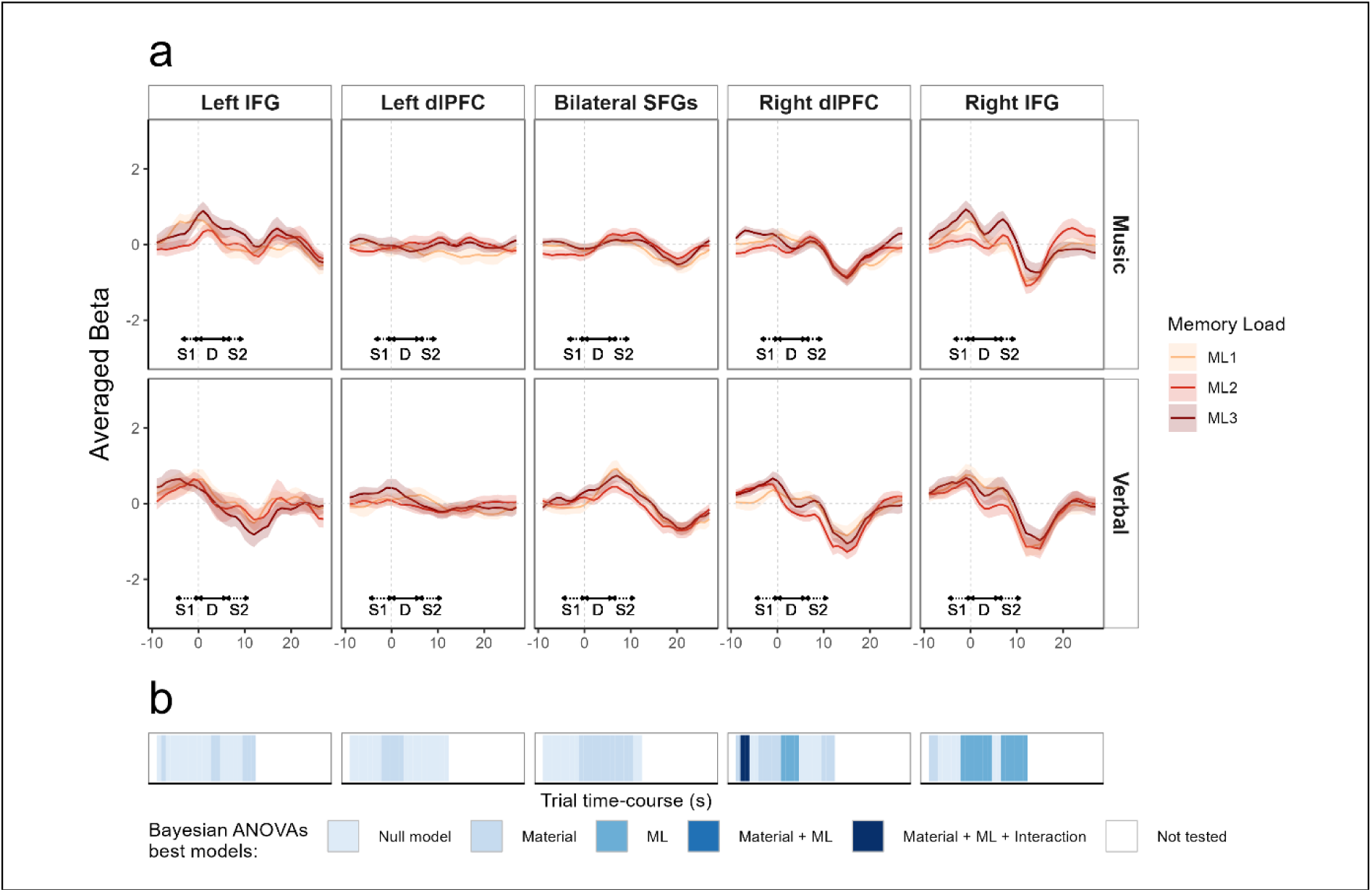
(a) Average beta (plain line) and standard error (shaded area) from the FIR deconvolution performed on HbR data for all participants (n=24), across five ROIs (left and right IFG, left and right dlPFC, bilateral SFGs), in a time window ranging from –9 to 27 seconds around delay onset (grey dotted vertical line), for the three memory load levels (ML1/ML2/ML3), for the musical material (top panel) and verbal material (bottom panel). For clarity purposes, S1, silent retention delay (D), and S2 durations are indicated with double-headed arrows, S1 and S2 arrows are dotted to indicate their variable duration according to the memory load. (b) Time-course representation of Bayesian analysis, each blue shade represents the best model as compared to the null model (BF_10_ > 1). No statistics were performed beyond 12 seconds after delay onset (end of S2 for the longest sequence + peak of the hemodynamic response, ∼ 6s).

